# Competing regulatory modules control the transition between mammalian gastrulation modes

**DOI:** 10.1101/2025.05.07.652670

**Authors:** Alexandra E. Wehmeyer, Johanna K. Schmitt, Felix Eggersdorfer, Lea Zissel, Chiara M. Schröder, Mehmet Tekman, André Dias, Katrin M. Schüle, Simone Probst, Alfonso Martinez-Arias, Katie McDole, Sebastian J. Arnold

## Abstract

During mammalian gastrulation cells of the primary germ layers are generated in anterior-to-posterior sequence employing different morphogenetic modes. Initial gastrulation is characterized by cell ingression through the early primitive streak, followed by posterior embryonic axis elongation via cell recruitment from progenitor pools. Molecular details of different genetic programs controlling early and late gastrulation remain ill described. Here, we employed stem cell-based mouse gastruloids to reveal two consecutively acting regulatory modules that orchestrate spatiotemporal progression of gastrulation. The early anterior module consists of the Tbx transcription factor *Eomes*, and signalling molecules *Nodal* and *Wnt3* that initiate gastrulation and generate anterior mesoderm and definitive endoderm from the early streak. The anterior module represses the second, *Tbxt/Wnt3a* posterior regulatory module controlling axial extension at trunk levels. Both circuitries are self-reinforcing while mutually repressing the counteracting module at multiple levels as the molecular basis for the spatiotemporal progression of gastrulation along the AP axis.

## Introduction

The onset of gastrulation in mouse is marked by the emergence of the primitive streak (PS), a transient structure formed by the ingression of epiblast cells that give rise to the nascent mesoderm cell layer (Arnold & Robertson, 2009). Initially, the PS forms as a confined hub of cells at the prospective posterior side of the cup-shaped embryo at embryonic day 6.5 (E6.5). From here the PS progressively elongates distally until E7.5, when the embryonic node forms at the most distal tip of the embryo (Downs & Davies, 1993; Williams et al., 2012). Early ingressing cells predominantly generate nascent mesoderm, including extraembryonic mesoderm (ExMes) and anterior, cranio-cardiac mesoderm (AM), followed by definitive endoderm (DE) progenitors that subsequently integrate into the outside endoderm layer (Kwon et al., 2008; Lawson et al., 1986; Probst et al., 2021; Viotti et al., 2014). After the first day of gastrulation, the morphogenetic mode to generate the progenitors that give rise to cells for the posterior extension of the body axis progressively changes (Aires et al., 2018; Bénazéraf & Pourquié, 2013; Duarte et al., 2023; Wymeersch et al., 2021). Different to epiblast cell ingression through the early PS, in later phases of gastrulation the generation of primordial tissues is fuelled from populations of proliferating progenitors. For example, progenitors with neural and mesodermal potential (NMPs) are initially found in regions of the node-streak border and the caudal lateral epiblast (Cambray & Wilson, 2007). NMPs persist as self-renewing progenitor pool in the chordo-neural hinge (CNH) of the posterior tail bud regions of embryos from where they contribute to the generation of mesoderm and spinal cord progenitors until gastrulation ceases (Cambray & Wilson, 2007; Wymeersch et al., 2019).

Gastrulation onset and the continuous patterning of the PS relies on instructive signals, including *Nodal/Smad2/3* and canonical Wnt signalling (Arnold & Robertson, 2009; Bardot & Hadjantonakis, 2020; Morgani & Hadjantonakis, 2020; Robertson, 2014). Epiblast expression of *Nodal* initiates the formation of the PS and instructs cell specification of early PS-derivatives, including *Mesp1*-expressing cranio-cardiac and extraembryonic mesoderm (Costello et al., 2011; Lu & Robertson, 2004), and anterior PS (APS) derived tissues, namely DE, node, notochord, and prechordal plate (Dunn et al., 2004; Vincent et al., 2003). PS formation additionally relies on canonical Wnt-signalling (Liu et al., 1999; Ten Berge et al., 2008). First, *Wnt3* is expressed at the site of the future PS and then along the PS until E7.5 (Liu et al., 1999; Rivera-Pérez & Magnuson, 2005)*. Wnt3* expression is followed by the upregulation of *Wnt3a* in the late PS and tail bud, that in concert with transcription factors (TFs) such as *Tbxt* (aka *Brachyury* or *T*) and *Cdx factors* promotes mesoderm lineages and the elongation of the posterior body axis (Amin et al., 2016; Aulehla et al., 2003; Herrmann et al., 1990; Martin & Kimelman, 2008; Savory et al., 2009; Takada et al., 1994; Young et al., 2009). This consecutive expression of different Wnt ligands during gastrulation suggests individual regulatory functions in the course of germ layer formation (Wang et al., 2012; Yamaguchi, 2008).

Cell type specification and germ layer morphogenesis additionally relies on sets of TFs, including the Tbx factors *Eomes* and *Tbxt*. *Eomes* instructs the specification of AM derivatives (Costello et al., 2011; Probst et al., 2021) and DE from the early PS (Arnold et al., 2008; Teo et al., 2011), and *Tbxt* is required for axial mesoderm derivatives, and paraxial mesoderm caudal to the first seven to ten pairs of somites (Koch et al., 2017; Schüle et al., 2023). In the absence of both Tbx TFs all mesoderm and endoderm (ME) gene programs entirely fail due to crucial requirements of Tbx factors for pluripotency exit, the generation of chromatin-based competence for ME lineage specification, and the concomitant repression of neuroectoderm lineage fates Previous studies suggest that *Eomes* is regulated by and functionally associated with *Nodal*/SMAD2/3-signalling (Brennan et al., 2001; Pfeiffer et al., 2018). Similarly, *Tbxt* is induced and functionally linked to canonical Wnt-signalling (Arnold et al., 2000; Martin & Kimelman, 2008; Yamaguchi et al., 1999). However, to date, the molecular interdependencies of signals and TF activities that orchestrate gastrulation along tight spatiotemporal patterns remain poorly understood.

Here, we employed 3D mouse embryonic stem cell-based gastruloids (Beccari et al., 2018; McNamara et al., 2024; Rossi et al., 2021; van den Brink et al., 2014) to untangle the dynamic interdependencies between signals and transcriptional regulators that govern the progression between different gastrulation stages as found in E6.5 to ∼E8.5 mouse embryos. To address individual functions of signalling components and TFs, we used variations of the classical gastruloid protocol (Beccari et al., 2018; Turner et al., 2017; van den Brink et al., 2014; Veenvliet et al., 2020) in combination with genetic approaches (Wehmeyer et al., 2022). Hence, we could define two mutually competing regulatory modules that contain the core molecular players essential for the progression through gastrulation stages. A rapid switch between these regulatory circuitries accounts for proper timing and coordination of gastrulation modes and coincides with the timepoint of formation of the node at E7.5 in the mouse embryo.

## Results

### Consecutive morphogenetic modes of gastrulation correspond to a transition of signalling signatures

Previous studies suggested that the cranial somites up to the level of somite pair 7-10 originate from early specified mesoderm and originate from a different progenitor pool than later forming, more caudal somites (Guibentif et al., 2021; Kinder et al., 1999). These progenitor pools most likely also underlie different molecular regulation. This notion is also supported by mouse mutants for *Tbxt* or *Wnt3a* that form most anterior somites but lack somites posterior to the first 7-10 pairs (Kispert & Herrmann, 1994; Takada et al., 1994). First, we aimed to inquire when different progenitor pools for anterior (<7 somites) and more posterior (>7 somites) mesoderm are generated. We therefore re-analysed previous live embryo timelapse cell tracking movies (McDole et al., 2018) to resolve when somitogenic mesoderm for somites at different levels is generated. Analysed embryos comprised the time span from early PS formation to the 8-10 somite stage, corresponding to E6.5 – E8.25 (McDole et al., 2018). We backtracked cells of the first 8 rows of cranial somites to the time of their origin (Fig. 1A). We found that the progenitors of most cranial somites are present in distal portions of the PS and are already generated before the embryonic node is formed (Fig. 1A, green cells, 11h40). Somitogenic mesoderm of more caudal somites is generated after the node becomes morphologically discernible (Fig. 1A, purple cells, 42h00), which corresponds to the timepoint E7.5 (Downs & Davies, 1993). Notably, the timepoint of E7.5 also indicates the onset of NMPs formation in vicinity to the embryonic node (Wymeersch et al., 2021), further supporting that the origin and mode of somitogenic mesoderm greatly differs between the most rostral and more caudal somites. To correlate the onset of a morphogenetic transition of gastrulation at E7.5 to changes in the signalling environment, we analysed published scRNA-seq data sets (Pijuan-Sala et al., 2019). We specifically examined the expression dynamics of known regulators of PS patterning, such as signalling molecules *Nodal*, *Wnt3* and *Wnt3a*, and the Tbx TFs *Eomes* and *Tbxt* (Fig. 1B). In accordance with previous expression analyses (Liu et al., 1999; Lu & Robertson, 2004; Takada et al., 1994), the scRNA-seq data similarly shows a rapid change of the signalling landscape around E7.5 (Fig. 1B). *Nodal* and *Wnt3* are lost abruptly around E7.5 from the PS, and in reverse *Wnt3a* expression is rapidly established and continues to be expressed in caudal mesoderm partially overlapping with *Tbxt* (Fig. 1B). *Eomes*, which is widely expressed throughout the posterior epiblast, PS and nascent mesoderm during early gastrulation (E7.0) and overlaps with *Tbxt* in the E7.0 and E7.5 PS, is similarly rapidly downregulated and mostly absent at E8.0 (Schüle et al., 2023). In sum, the highly dynamic expression patterns of these key regulators of gastrulation tightly correspond to the transition of gastrulation stages at the time of embryonic node formation at E7.5.

**Fig. 1.**
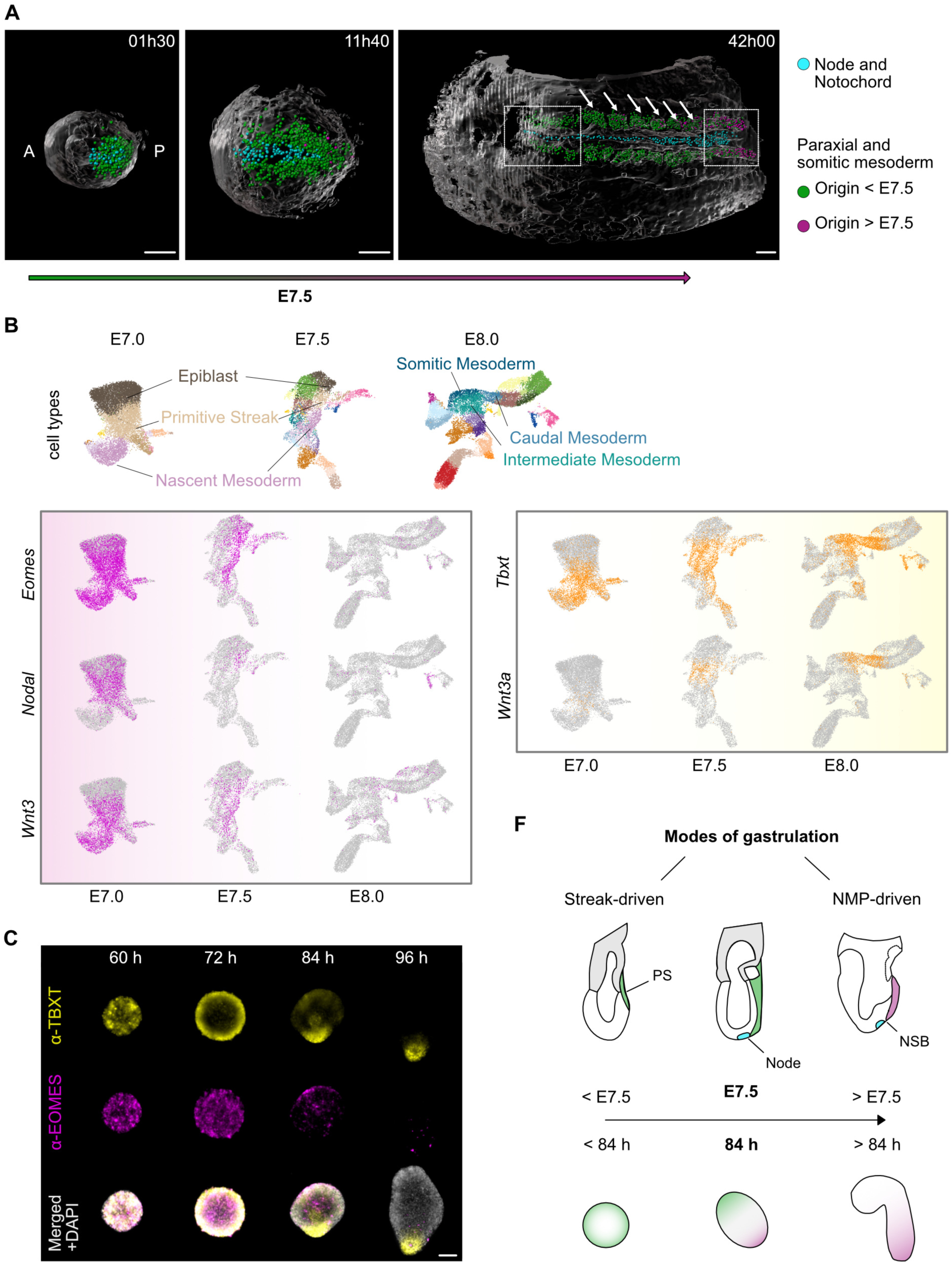
Morphogenetic gastrulation modes and signalling environments switch at E7.5. **(A)** Snapshot images showing cell tracking from beginning of gastrulation (01h30) to the time of node formation (11h40) until 9-somite stage (42h00) (Ventral views, full movie as Supplemental movie 1). Cells of newly forming somitogenic paraxial mesoderm are marked in green and purple depending to the time of origin, either prior to node formation (<E.7.5, green cells), or post node formation (>E.7.5, purple cells). Blue cells mark node and notochord. At 42h00 somites are indicated by arrows. The first two somite-pairs are not visible as epithelial structures (boxed area left), and unsegmented paraxial mesoderm is indicated as boxed area (boxed area right). The rostral 8 somites are almost exclusively formed from presomitic mesoderm, that was already formed prior to E7.5, while later emerging mesoderm (purple) contributes to more caudal somites. Scale bars 100 µm. **(B)** Gene expression signatures around E7.5 shown as UMAPs of published single cell mouse embryo RNAseq data sets (Pijuan-Sala et al., 2019). *Eomes* and *Tbxt* are co-expressed in subpopulations of the PS and mesoderm prior to the appearance of the node at E7.5, overlapping with *Nodal* and *Wnt3* expression. At E7.5, *Eomes, Nodal* and *Wnt3* are downregulated. After E7.5, emerging mesoderm is established in the presence of *Tbxt* and *Wnt3a*. Extraembryonic tissues are excluded from this UMAP representation. **(C)** Immunofluorescent staining (IF) of CHIR-treated wildtype (WT) mouse gastruloids. Dynamic changes in TBXT and EOMES protein distribution are shown in 12 h intervals. Similar to embryonic expression, TBXT and EOMES are initially co-staining (60 h) before their domains separate. At 84 h EOMES is downregulated and TBXT polarizes to one pole, from where gastruloid elongation occurs. n≥5. Scale bars 100 µm. **(D)** Schematic of gastrulation stages in embryos and the corresponding timepoints in mESC-derived gastruloids. Before E7.5, nascent mesoderm is generated by cell ingression through the PS which is recapitulated in gastruloids <84 h, when EOMES and TBXT are co-localizing. Around E7.5 the node forms at the distal tip of the embryo. *Eomes, Nodal* and *Wnt3* are rapidly downregulated and neuro-mesodermal progenitors (NMPs) form in vicinity of the node. Similar regulation is found at 84 h in gastruloids. >E7.5 and >84 h, posterior elongation of the embryo and of gastruloids is accomplished in an environment of high WNT3a and TBXT levels. Primitive streak (PS), Node streak border (NSB), Neuro-mesodermal progenitor (NMP).

### Gastruloids recapitulate the transition of gastrulation modes

To study the progression between gastrulation modes from early PS stages to posterior axial elongation after E7.5, we employed embryonic stem cell (ESC)-based gastruloids. We first applied the standard protocol of gastruloid formation using a pulse of CHIR-treatment to induce canonical Wnt-signalling responses (van den Brink et al., 2014) (Fig. S1A) and analysed the protein dynamics of EOMES and TBXT as key regulators of ME formation, by immunofluorescence (IF) staining (Fig. 1C). Similar to the early embryo (Schüle et al., 2023), EOMES and TBXT initially co-localise in 60h gastruloids before TBXT becomes restricted to outer cells of gastruloids at 72 h. At 84 h obvious morphological asymmetries become apparent when TBXT is found at one pole of the gastruloids, and EOMES is reduced and sparsely found at the opposite pole. Elongation of gastruloids follows the loss of EOMES at 96 h, thereby mimicking embryonic development after node formation (E7.5), when axial elongation is governed by NMP-generated tissues. Hence, patterns of expression dynamics of EOMES and TBXT in gastruloids largely reflect the observations at the embryonic streak. Accordingly, gastruloids can be used as suitable model systems for studies of the mechanisms that control the transition between gastrulation stages from <E7.5 to >E7.5 (Fig. 1D).

### Combined activities of Tbx factors and signals instruct temporal progression and patterning of gastruloids

To dissect the molecular regulation of gastrulation progression from early PS stages to posterior axis elongation we first investigated functional requirements for *Eomes* and *Tbxt* during gastruloid formation in context of different signalling environments. We used either a standard CHIR pulse, or induced with the NODAL-analogue ACTIVINA (ActA) (Fig. S1A, B). We compared gastruloids from wildtype (WT), *Eomes*- or *Tbxt-* deficient (*Eo ^−/−^* or *Tbxt ^−/−^)* and double knockout (dKO) mESCs (Fig. S1C) (Schüle et al., 2023; Tosic et al., 2019). Expectedly, using the standard CHIR protocol, WT gastruloids stereotypically develop a cell-dense region, representing the anterior pole, and elongate posteriorly (Fig. 2A, S1D). The simultaneous deletion of both *Eomes* and *Tbxt (*dKO) fully abrogates symmetry breaking, and dKO aggregates remain round (Fig. S1D). *Tbxt* ^−/−^ gastruloids fail to elongate and remain oval-shaped at 120 h, as also previously reported (Fig. 2B, S1D) (Wehmeyer et al., 2022). Surprisingly, *Eo ^−/−^* gastruloids elongate and are morphologically similar to WT (Fig. 2C; Fig. S1D), despite crucial morphogenetic roles of *Eomes* during early gastrulation (Arnold et al., 2008). We morphometrically quantified these observations by measuring the length and width of gastruloids and calculated length/width ratios for each genotype and condition as an objective read-out for elongation (Fig. S1E, F, Supplementary Table 1).

**Fig. 2.**
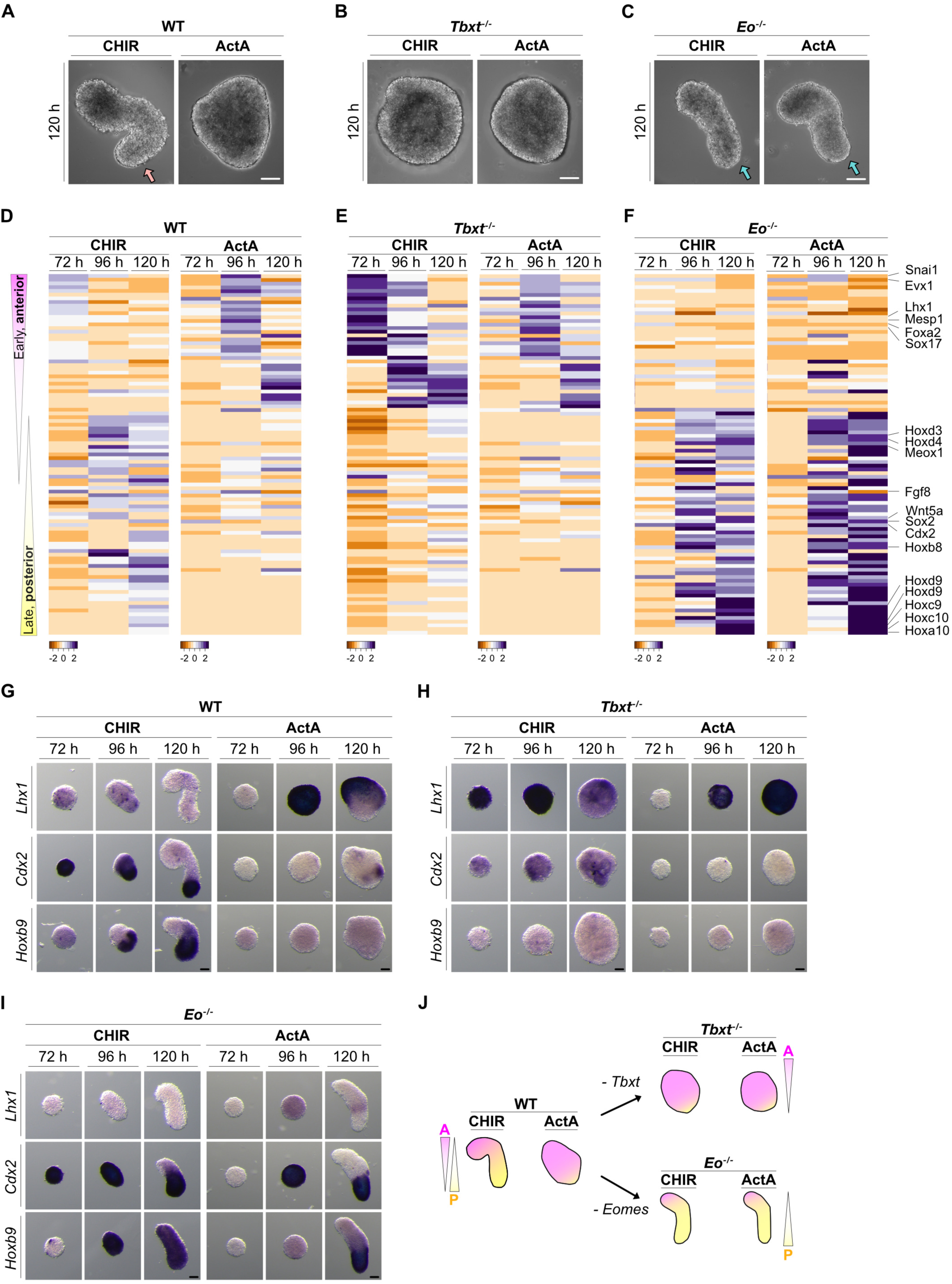
Anterior-posterior axis elongation in gastruloids relies on Tbx TFs to transduce instructive signals. **(A-C)** Brightfield images of representative gastruloids at 120 h from **(A)** wildtype (WT), **(B)** *Tbxt ^−/−^* or **(C)** *Eomes*-deficient (*Eo* ^−/−^) mouse embryonic stem cells (mESCs) that were generated by pulsed induction (48 h −72 h) with CHIR, or ActivinA (ActA). Arrows indicate posterior elongation. n≥60. Scale bars 100 µm. Compare Fig. S1 for additional phenotypic analysis. **(D-F)** Molecular profiling by RNAseq. Heatmap depicts spatial and temporal expression changes of characteristic anterior and posterior marker genes (n=97, Supplementary Table 2,3). **(D)** CHIR-treated WT gastruloids indicate the sequence of early, anterior marker gene expression and the transition to later expressed posterior marker genes. ActA-treated WT gastruloids shows a delayed emergence of anterior markers and strongly reduced posterior marker genes. **(E)** *Tbxt ^−/−^* gastruloids show an upregulation of anterior, and downregulation of posterior marker genes. **(F)** *Eo* ^−/−^ gastruloids exhibit the opposite pattern: an upregulation of posterior and decreased expression of anterior marker genes. Remarkably, the gross anterior-posterior patterning of *Tbxt ^−/−^* and *Eo* ^−/−^ gastruloids is largely independent of the upstream pulse being either CHIR or ActA. **(G-I)** Whole-mount *in situ* hybridization (WISH) of an anterior marker (*Lhx1*), and posterior marker genes (*Cdx2, Hoxb9*) illustrates anterior-posterior pattering at 24 h intervals of CHIR or ActA pulsed gastruloids of indicted genotypes, **(G)** WT, **(H)** *Tbxt* ^−/−^ and **(I)** *Eo* ^−/−^. Marker genes reflect the patterning phenotypes deduced from heatmap analysis. n≥6. Scale bars 100 µm. **(J)** Schematic summarizing anterior-posterior patterning of WT, *Tbxt ^−/−^* and *Eo* ^−/−^ gastruloids in different signalling environments by stimulation with either CHIR, or ActA. A - anterior. P - posterior.

Next, we phenotypically analysed gastruloids, when induced with ActA (25 ng/ ml), that also was applied as pulse between 48 and 72 h (Fig. S1B). As previously reported (Turner et al., 2017; van den Brink et al., 2014), ActA suppresses axis elongation in WT aggregates, which acquire irregular round to triangular shapes at 120 h similar to *Tbxt ^−/−^* aggregates (Fig. 2A, S1D). Unexpectedly, *Eo ^−/−^* (ActA)-induced gastruloids consistently elongate and are morphologically alike CHIR-treated gastruloids. These findings suggest that *Tbxt* promotes axis-elongation in conjunction with CHIR/Wnt-stimulation, and that *Eomes* mediates elongation-inhibiting activities of the Nodal/ActivinA-signalling pathway.

To molecularly characterize these morphological findings in gastruloids we transcriptionally profiled CHIR- and ActA-induced gastruloids from WT, *Tbxt ^−/−^* and *Eo ^−/−^* cells at three time point (72 h, 96 h and 120 h) using bulk RNA sequencing and analysed differentially expressed genes (DEGs) when compared to WT gastruloids (Fig. 2D, E, F, S2A-D). Gross numbers of DEGs indicate that the deletion of *Tbxt* results in larger transcriptional differences in CHIR treated gastruloids (Fig. S2A), and the deletion of *Eomes* leads to largest effects in ActA induced gastruloids (Fig. S2B), suggesting functional synergism between pairs of *Tbxt*/Wnt-(CHIR)-signalling and *Eomes*/Nodal/ActA-signalling, respectively. GO Term analyses of DEGs confirm the phenotypic observations of disturbed AP axis formation and patterning of *Tbxt ^−/−^* (CHIR)-induced gastruloids, that is remarkably reestablished in ActA-treated aggregates when *Eomes* is deleted (*Eo ^−/−^* (ActA); Fig. S2C, D).

To illustrate the transcriptional patterns characteristic for gastruloid development, we generated heatmaps that show the sequential expression of marker genes along the anterior-to-posterior axis (n=97 genes, Supplementary Tables 2, 3). The heatmap for WT, CHIR-treated gastruloids shows the orderly pattern of early arising (72 h) anterior marker genes (e.g., *Snail, Evx1, Lhx1, Mesp1*) and the transition to later expressed marker genes of posterior identities (e.g., *Cdx2, Wnt5a, Hoxa9, Hoxc10*) (Fig. 2D, CHIR-treatment). Heatmaps across different induction regimes and genotypes show gross alterations of spatiotemporal patterns. Pulsing WT gastruloids with ActA attenuates expression of posterior and increases expression of anterior markers (Fig. 2D). Additionally, ActA leads to the globally delayed expression of early anterior marker genes from 72 to 96 h in WT and *Tbxt*-deficient gastruloids, supposedly reflecting the pluripotency-maintaining effect of ActA/Nodal-signalling (Vallier et al., 2009) (Fig. 2D, E). *Tbxt ^−/−^* gastruloids show markedly increased expression of anterior and severely reduced expression of posterior marker genes, irrespective of pulsing with either CHIR or ActA (Fig. 2E). In reverse, *Eomes*-deficient (*Eo ^−/−^)* gastruloids show a gross reduction of early anterior and increased expression of posterior markers above levels found in WT gastruloids, irrespective of the initial pulse of CHIR or ActA (Fig. 2F, compare to Fig. 2D). Importantly, despite ActA-treatment *Eo ^−/−^* gastruloids strongly express markers of posterior axis identity at 120 h fitting to the observed elongated phenotype (Fig. 2C). The heatmap analyses of AP-axis markers from RNA-seq data were confirmed by whole-mount in situ hybridisation (WISH) of early anterior (*Lhx1*) and later, posterior (*Cdx2*, *Hoxb9*) expressed marker genes (Fig. 2G-I). WISH marker analysis validated that *Nodal*/ActA promotes anterior fate, while Wnt/CHIR-signals instruct posterior identities (Fig. 2G-I). These patterning functions of signals critically rely on the Tbx factors *Eomes* and *Tbxt*. Irrespective of the initial inductive signalling pulse, gastruloids acquire posterior identities and elongate in the absence of *Eomes* (Fig. 2F, I), while *Tbxt*-deficient gastruloids acquire mainly anterior fate identities (Fig. 2E, H). This suggests that activities of *Nodal*- and Wnt-signalling in gastruloids are predominantly mediated by the presence or absence of *Eomes* or *Tbxt*, respectively (Fig. 2J).

### EOMES and BRACHYURY show mutual repressive regulation

Anterior-posterior marker expression in gastruloids largely relies on opposing activities of *Eomes* and *Tbxt* (Fig. 2). Competing activities are also reflected in differential protein distribution within gastruloids (Fig 1D). To capture the time period when the AP-asymmetry is established, we performed IF staining for EOMES and TBXT in CHIR-treated WT gastruloids collected from the start of CHIR treatment at 48 h in 6 h intervals until 96 h (Suppinger et al., 2023) (Fig. 3A, partially also shown in Fig. 1C). EOMES and TBXT initially co-localise from 60 h until 72 h, before becoming restricted to different domains at 84 h (Fig. 3A). To address if the expression of both Tbx factors impact each other, we compared IF pattern found in WT gastruloids to *Eo* ^−/−^ and *Tbxt ^−/−^*gastruloids (Fig. 3B, C). In *Eo ^−/−^* gastruloids, polarization of TBXT to one side of the gastruloid takes place, however, overall protein distribution is less confined compared to WT gastruloids (Fig. 3B). In *Tbxt ^−/−^* gastruloids, EOMES is first induced similar to WT, but in contrast to the downregulation in WT, increased EOMES staining persists at 96 h in *Tbxt ^−/−^* (Fig. 3C). To relate IF staining to gene expression levels we quantified expression from RNAseq data at 72, 96 and 120 h (Fig. 3D, E). mRNA levels grossly match the observed protein staining intensities, such that we find increased *Tbxt* levels in *Eo ^−/−^* gastruloids (Fig. 3E), and maintained *Eomes* expression in *Tbxt ^−/−^* gastruloids (Fig. 3D). These findings suggest a mutual negative regulation after an early phase of co-expression by *Eomes* and *Tbxt* as schematically depicted (Fig. 3F).

**Fig. 3.**
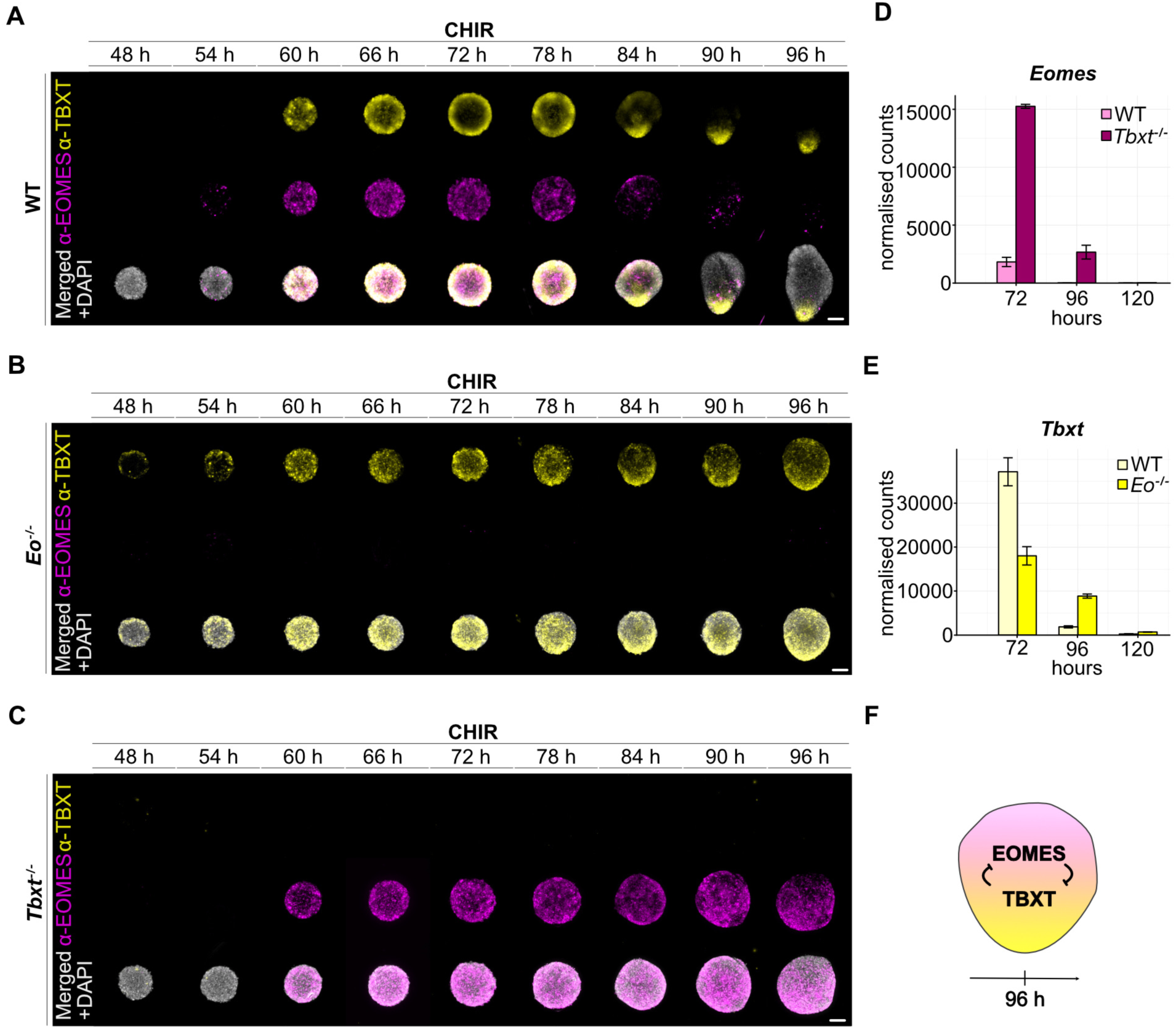
EOMES and TBXT are dynamically distributed and mutually repress each other. **(A-C)** Immunofluorescent staining of CHIR-treated WT, *Eo* ^−/−^ and *Tbxt ^−/−^* gastruloids in 6 h intervals to demonstrate dynamic changes of TBXT and EOMES during the early phase (48 - 96 h) of asymmetry breaking along the anterior-posterior axis. **(A)** TBXT and EOMES initially co-localise (60 h) in WT gastruloids before domains separate (72 h). At 96 h EOMES is largely absent and TBXT polarizes to one pole. **(B)** *Eo* ^−/−^ gastruloids exhibit premature TBXT at slightly increased levels. TBXT is less clearly polarized at 96 h compared to WT gastruloids. **(C)** *Tbxt ^−/−^ gastruloids* initiate EOMES similar to WT and maintain high levels at later timepoints (96 h), in contrast to WT that loose EOMES from 90 – 96 h. n≥5. Scale bars 100 µm. **(D, E)** RNA levels of *Tbxt* and *Eomes* in CHIR-treated WT, and *Eo* ^−/−^ or *Tbxt ^−/−^* gastruloids reflect the protein dynamics. **(D)** At 96 h, *Tbxt* mRNA levels are increased in *Eo* ^−/−^, and **(E)** *Eomes* levels are maintained longer in *Tbxt ^−/−^* gastruloids until 96 h. Protein and mRNA expression dynamics suggest mutual negative regulation between *Eomes* and *Tbxt* as **(F)** summarized in a schematic. RNA expression levels are plotted as normalised counts of three replicates. Error bars indicate SEM.

### Tbx TFs and signalling ligands act as reinforcing functional modules of Eomes/Nodal/Wnt3 *and* Brachyury/Wnt3a

Next, we aimed to investigate more deeply potential modes of mutual repression between *Eomes* and *Tbxt*. Based on the delayed repression dynamics (Fig. 3) we suspected cell non-cell-autonomous reciprocal repression mechanisms between EOMES and TBXT in addition to previously reported direct repression of *Tbxt-*activities by *Eomes* (Schüle et al., 2023). Hence, we analysed RNA expression patterns of the upstream signalling regulators of *Eomes* and *Tbxt* expression, namely *Nodal* (Brennan et al., 2001; Simon et al., 2017), *Wnt3* (Pfeiffer et al., 2018), and *Wnt3a* (Arnold et al., 2000; Yamaguchi et al., 1999), using WISH at 48 h, 72 h, 84 h, 96 h, and 120 h in WT, *Tbxt ^−/−^* and *Eo* ^−/−^ gastruloids (Fig. 4A-C). In WT gastruloids *Nodal* expression precedes the CHIR pulse at 48 h, expression peaks at 72 h, and is followed by downregulation in most parts of the gastruloids until 96 h (Fig. 4A). This pattern is reminiscent of the embryonic expression where *Nodal* is present in the epiblast before PS formation and becomes confined to the streak followed by the rapid downregulated at E7.5 (Collignon et al., 1996). In WT gastruloids *Wnt3* and *Wnt3a* are first expressed after the CHIR pulse post 72 h. *Wnt3* is downregulated between 72 h and 96 h, while *Wnt3a* remains highly expressed and becomes confined to the posterior tail-like region, that coexpresses *Tbxt* (Wehmeyer et al., 2022) (Fig. 4A). Hence, also *Wnt3* and *Wnt3a* expression reflect the kinetics found in embryos where *Wnt3* expression is lost from the PS at E7.5 (Liu et al., 1999), and *Wnt3a* expression is maintained in the tail bud (Takada et al., 1994).

**Fig. 4.**
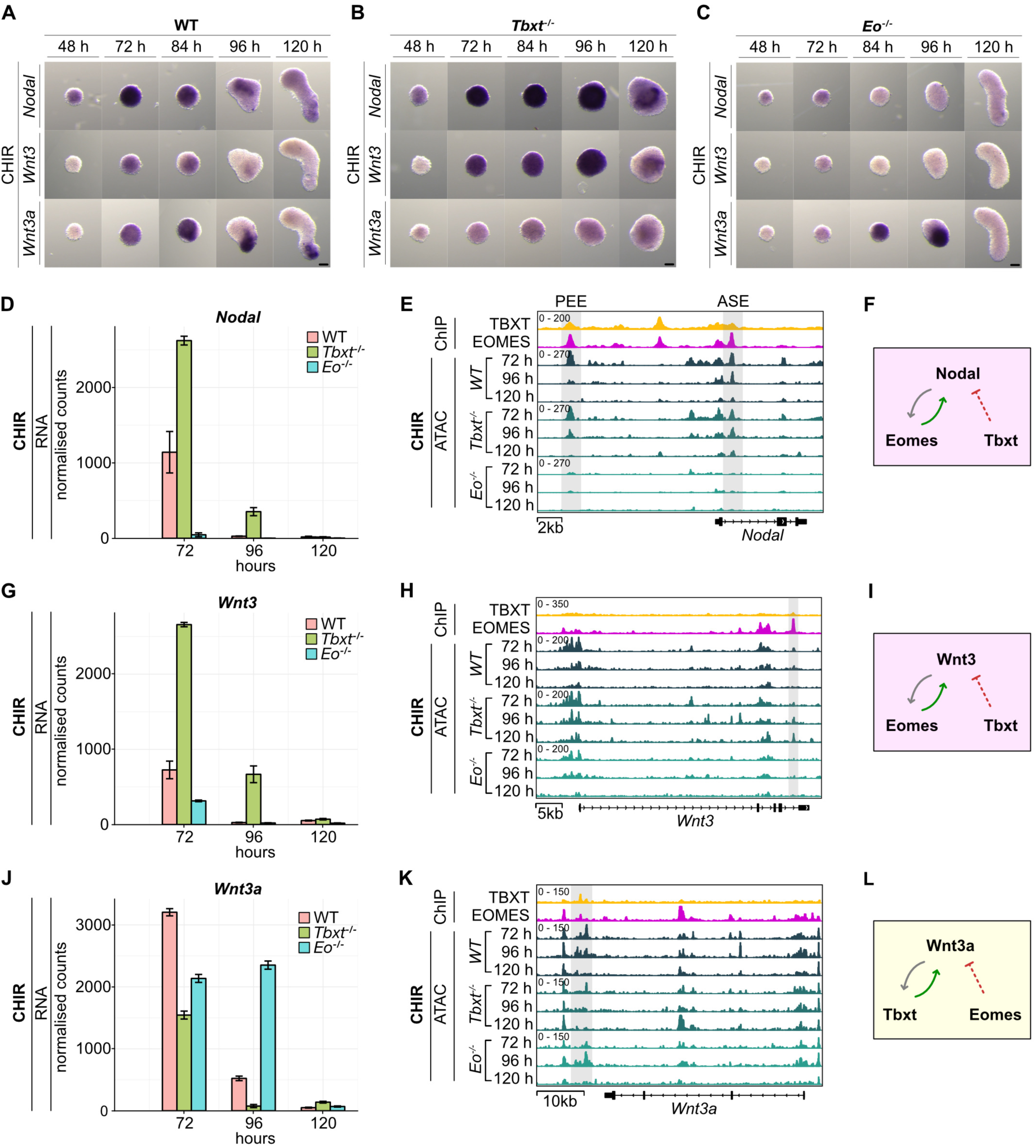
Tbx factors and signals act as self-reinforcing functional modules of Eomes/Nodal/Wnt3 *and* Tbxt/Wnt3a. **(A-C)** Expression analysis of main upstream signalling regulators of *Eomes* (*Nodal*, *Wnt3)* and *Tbxt* (*Wnt3a)* by WISH in WT, *Tbxt* ^−/−^ and *Eo* ^−/−^ CHIR-pulsed gastruloids **(A)** In WT, *Nodal* expression precedes the exogenous CHIR-pulse at 48 h, expression peaks at 72 h, and is subsequently downregulated. *Wnt3* and *Wnt3a* are expressed after the CHIR pulse from 72 h. *Wnt3* is downregulated between 72 and 96 h, while *Wnt3a* remains expressed in the posterior tail-bud like region. **(B)** *Tbxt* ^−/−^ gastruloids, show increased expression of *Nodal* and *Wnt3* that are maintained beyond 96 h, while they are devoid of *Wnt3a* expression. **(C)** *Eo ^−/−^* gastruloids show opposing expression patterns and lack *Nodal* and *Wnt3* expression, while strongly expressing polarized *Wnt3a* at 96 h. n≥6. Scale bars 100 µm. **(D-L)** Analysis of the regulation and interdependency of signals and Tbx TFs. mRNA expression of **(D)** *Nodal*, **(G)** *Wnt3* and **(J)** *Wnt3a* in WT, *Tbxt* ^−/−^ and *Eo* ^−/−^ CHIR-pulsed gastruloids recapitulate observations of WISH analysis. **(E, H, K)** ChIPseq and ATACseq coverage tracks at the gene loci of *Nodal, Wnt3 and Wnt3a* show ChIP-binding of TBXT and EOMES to putative enhancer sites (indicated in grey) that undergo dynamic changes of chromatin accessibility by ATACseq. **(E, H)** Accessibility at the *Nodal* and *Wnt3* locus is reduced in the absence of *Eomes*. **(K)** The *Wnt3a* locus shows reduced accessibility in *Tbxt* ^−/−^ gastruloids. Counts normalized to RPKM are indicated in E, H, K. **(F, I, L)** Schematics summarizing the suggested cross-regulation of signals and Tbx TFs.

In *Tbxt ^−/−^*gastruloids *Nodal* and *Wnt3* expression levels are markedly increased and maintained beyond 96 h, while *Wnt3a* expression is almost absent (Fig. 4B). In contrast, *Eo ^−/−^* gastruloids show the opposite expression dynamics of decreased *Nodal* and *Wnt3*, while *Wnt3a* levels appear grossly normal and localise to a single, enlarged expression domain at the posterior pole at 96 h (Fig. 4C). These expression patterns by WISH are also recapitulated in mRNA levels of RNAseq data (Fig. 4D, G. J). We next asked if expression of *Nodal*, *Wnt3* and *Wnt3a* are directly regulated by EOMES and/or TBXT. Hence, we analysed chromatin binding of TBXT and EOMES by ChIPseq (chromatin immunoprecipitation coupled with sequencing, data from Schüle et al., 2023). Here, we find binding of EOMES, TBXT or both to the genomic loci of *Nodal, Wnt3* and *Wnt3a* (Fig. 4E, H, K). To study the functional relevance of Tbx TF binding we performed ATACseq (Assay for Transposase-Accessible Chromatin with sequencing) in gastruloids as readout for Tbx factor activity (Schröder et al., 2025). At the *Nodal* locus, EOMES, and to minor degree also TBXT, bind the well-described proximal enhancer element (PEE) ∼12 kb upstream of the promoter that controls *Nodal* expression in the epiblast and the PS (Fig. 4E) (Norris & Robertson, 1999; Vincent et al., 2003) as well as to asymmetric element (ASE) enhancer of intron 1 (Norris et al., 2002). Chromatin accessibility at the PEE and the ASE is severely reduced in *Eo ^−/−^*gastruloids, suggesting direct regulation by EOMES (Fig. 4E), while accessibility is unaltered in *Tbxt ^−/−^*. EOMES also binds to the *Wnt3* locus, where *Eomes*-deletion leads to reduced chromatin accessibility at a putative intronic enhancer (Fig. 4H). These findings suggest direct positive regulation of *Nodal* and *Wnt3* by *Eomes* (Fig. 4F, I), but not by *Tbxt*. In reverse, at the *Wnt3a* locus we find reduced chromatin accessibility in *Tbxt ^−/−^*gastruloids at TBXT bound site (Fig. 4K) which could account for the reduced *Wnt3a* expression in *Tbxt* ^−/−^ gastruloids (Fig. 4L), as also previously suggested in zebrafish (Martin & Kimelman, 2008).

In summary, these data suggest that *Eomes* acts by a self-reinforcing feed-forward loop with the upstream regulators *Nodal* and *Wnt3* that together build a functional module that controls cell lineage specification and axial identity of early formed anterior cell types. In contrast, *Tbxt* forms an autoregulatory feed-forward loop with the upstream signalling regulator *Wnt3a*, both being crucially required for posterior axis elongation. Together the two functional modules of *Eomes/Nodal/Wnt3* and *Tbxt/Wnt3a* orchestrate patterning along the AP-axis of the PS.

### *Eomes* induces anterior fates in gastruloids by repression of *Tbxt* functions

Our previous experiments demonstrated that ActA treatment of WT gastruloids abrogates gastruloid elongation and expression of posterior markers in the presence of *Eomes* (Fig. 2A, D, G), but not in *Eomes ^−/−^* gastruloids (Fig. 2C, F, I, illustrated in Fig. 3J). We performed simultaneous IF for EOMES and TBXT in ActA-induced WT and *Eo ^−/−^* gastruloids (Fig. 5A) to test if the inhibition of axis-elongation by ActA results from the absence of TBXT in WT gastruloids. However, in WT(ActA) gastruloids TBXT is abundant and also shows the confined localization to one pole of the gastruloid at 120 h, albeit at reduced staining intensities when compared to CHIR induction (Fig. 5A, compare to Fig. 3A, E). Expectedly, ActA-pulsed WT gastruloids show high EOMES levels (Fig. 5A) that remain elevated at timepoints past 84 h, when EOMES is downregulated in WT(CHIR) gastruloids (Fig. 5A, compare Fig. 3A). ActA induced *Eo ^−/−^* gastruloids undergo axis elongation similar to CHIR-treated WT gastruloids (Fig. 3A). In accordance, TBXT staining in *Eo* ^−/−^(ActA) gastruloids remarkably resembles patterns of WT(CHIR) gastruloids, where TBXT first shows circumferential staining that later localizes to the posterior, extending tail bud-like region (Fig. 5A, compare to 3A). Thus, differences of TBXT patterns or levels are unlikely to accounts for the gross phenotypic differences between ActA treated WT and *Eo* ^−/−^ gastruloids. Therefore, we further investigated the mechanisms by which EOMES suppresses gastruloid elongation. We previously reported on competing activities of *Eomes* and *Tbxt* at the level of newly forming chromatin accessibility in differentiating mESCs (Schüle et al., 2023). In the current study, we thus tested if the forced expression of *Eomes* can also repress *Tbxt* functions required for axis elongation of gastruloids after the CHIR pulse. Remarkably, the doxycycline-induced expression of *Eomes* (Wehmeyer et al., 2022) simultaneous to the CHIR pulse (Fig. 5B, C) fully abrogates elongation of gastruloids (Fig. 5D, E). Inhibition of elongation is not due to absence of TBXT, since we found presence of TBXT after EOMES-induction by IF staining at 72 h with similar distribution but at slightly reduced intensities compared to WT gastruloids (Fig. 5F). From 96 h, TBXT is less confined and more dispersed after forced *Eomes*-expression, and at 120 h it is absent from the extending posterior pole. Mildly reduced levels of *Tbxt* mRNA are also seen in RNAseq analysis of gastruloids after forced *Eomes*-expression (Fig. 5G). The heatmap analysis of anterior-posterior genes in *Eomes*-induced gastruloids shows a transcriptional pattern that is strongly biased towards early expressed anterior marker genes (Fig. 5H, Supplementary Table 4). This pattern is similarly found in ActA-pulsed WT gastruloids, and in *Tbxt* ^−/−^ CHIR-pulsed gastruloids (compare with Fig. 2D, E). Thus, we next assessed if the lack of posterior marker expression after forced *Eomes* expression is due to compromised TBXT transcriptional functions, and performed WISH for *Tbxt* target genes, *Rspo3* and *Msgn1* (Evans et al., 2012; Schüle et al., 2023). Remarkably, EOMES expression drastically reduces expression of both *Rspo1* and *Msgn1* at 96 h and 120 h (Fig. 5I). This confirms previous observations of repressed *Tbxt*-functions by *Eomes* in embryoid bodies by currently poorly understood mechanisms (Schüle et al., 2023).

**Fig. 5.**
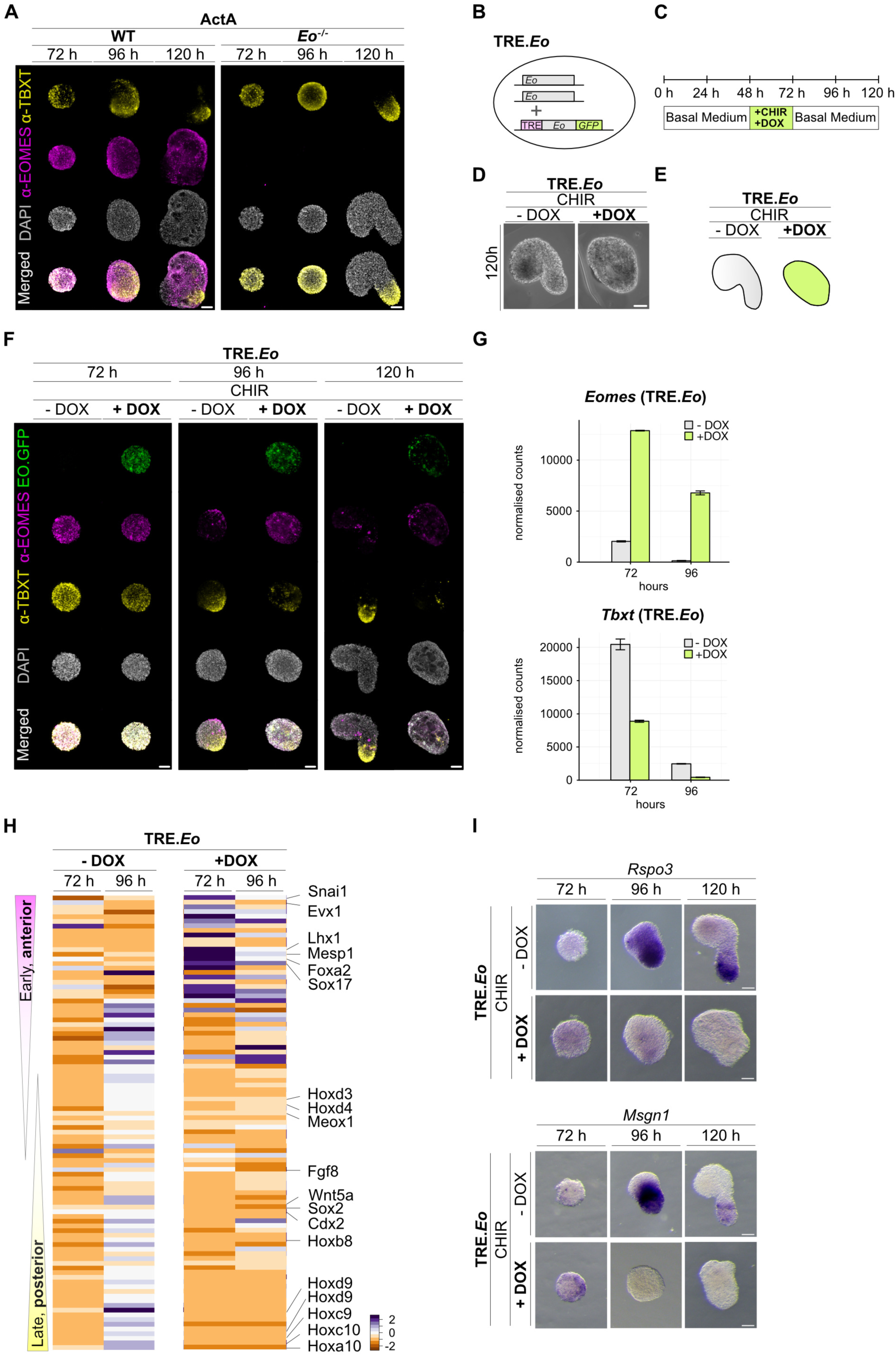
Repressive functions of ActivinA on axis elongation are mediated by EOMES. **(A)** IF staining of ActA-induced WT and *Eo* ^−/−^ gastruloids shows highly abundant EOMES in WT that is maintained until 120 h. TBXT is present and localized to one pole in WT and *Eo* ^−/−^ gastruloids, but posterior elongation is only promoted in absence of EOMES *(Eo* ^−/−^). n≥5. Scale bars 100 µm. **(B)** Schematic of the dox-inducible cell line (TRE.*Eo*) for the forced expression of *Eomes* in an otherwise wildtype *Eomes* background, and **(C)** schematic of the protocol used to generate CHIR-pulsed gastruloids with forced *Eomes*.GFP expression (CHIR+DOX). Control gastruloids are treated with CHIR only. **(D)** Brightfield images at 120 h demonstrate that forced *Eomes* expression impairs axis elongation, even in CHIR-induced gastruloids as schematically illustrated in **(E)**. **(F)** IF staining for EOMES, GFP and TBXT after forced *Eomes*.GFP expression in CHIR-pulsed gastruloids at indicated timepoints shows abundant TBXT at 72 and 96 h in uninduced and *Eomes*-induced cells, but absence of TBXT at 120 h after forced *Eomes* expression. n≥10. Scale bars 100 µm. **(G)** mRNA expression levels of *Eomes* and *Tbxt* at 72 and 96 h grossly recapitulate IF staining in TRE.*Eo* gastruloids (+DOX vs. -DOX) and show reduced and prematurely downregulated levels of *Tbxt*. **(H)** Heatmaps of RNAseq analysis of CHIR-pulsed gastruloids at 72 and 96 h shows the absence of posterior marker gene expression and increased anterior signature of gastruloids after forced *Eomes* expression (+Dox). Control gastruloids (-DOX) show the same expression dynamics as WT gastruloids (Fig. 2D). **(I)** WISH of two putative *Tbxt*-dependent target genes (*Rspo3* and *Msgn1*) with posterior expression in WT gastruloids confirms the absence of the posterior expression signature after forced *Eomes*-expression in gastruloids at 72, 96 and 120 h. n≥5. Scale bars 100 µm.

### *Eomes* represses canonical *Wnt3a* signalling at multiple levels

*Tbxt* and canonical Wnt-signalling act synergistically during posterior axial elongation (Arnold et al., 2000; Yamaguchi et al., 1999). Thus, we reasoned that repression of *Tbxt* functions might be additionally relayed through the suppression of the canonical Wnt-cascade by *Eomes*. First, we analysed expression of *Wnt3a*, normally acting in a positive feed-forward loop with *Tbxt,* in ActA-induced gastruloids in the presence (WT) and absence of *Eomes* (*Eo ^−/−^*) by fluorescent *in situ* hybridization (FISH) (Fig. 6A). While *Wnt3a* expression is largely absent from ActA-induced WT gastruloids, we find *Wnt3a* mRNA at the posterior pole of *Eo ^−/−^*(ActA) gastruloids at 120 h (Fig. 6A), and by analysis of RNAseq data at 96 and 120 h (Fig. 6B). The analysis of ATACseq data of WT and *Eo ^−/−^*gastruloids reveals a putative regulatory region 3’ adjacent to the *Wnt3a* gene locus (Fig. 6C) that is bound by EOMES and TBXT (Fig. 4K). The dynamic changes of chromatin accessibility at this region closely correlate to expression dynamics of *Wnt3a* observed in different conditions, and at different timepoints (Fig. 6C, D), suggesting that EOMES and TBXT directly contribute to the control of *Wnt3a* expression in an antagonistic manner. To assess more broadly canonical Wnt-signalling activities in gastruloids, we compared heatmaps of RNAseq data representing Wnt pathway components and the signalling signature (Fig. 6D). Here, we find a signature of Wnt pathway activation (*Axin2, Tcf7, Lef1* levels) in CHIR-treated WT, and in ActA-treated *Eo ^−/−^* gastruloids, but only largely reduced signature expression in ActA-treated WT gastruloids. The positive signature of canonical Wnt pathway activation in *Eo ^−/−^*gastruloids is accompanied by the reduced expression of inhibitory regulators of the Wnt pathway such as *Tcf7l1* and *Dkk1* (Fig. S3A, B). We also performed heatmap analysis of CHIR-pulsed gastruloids, that show an active Wnt signalling signature in WT and *Eo ^−/−^*gastruloids, that is broadly reduced in *Tbxt* ^−/−^ gastruloids in accordance with the observed phenotype (Fig. S3C).

**Fig. 6.**
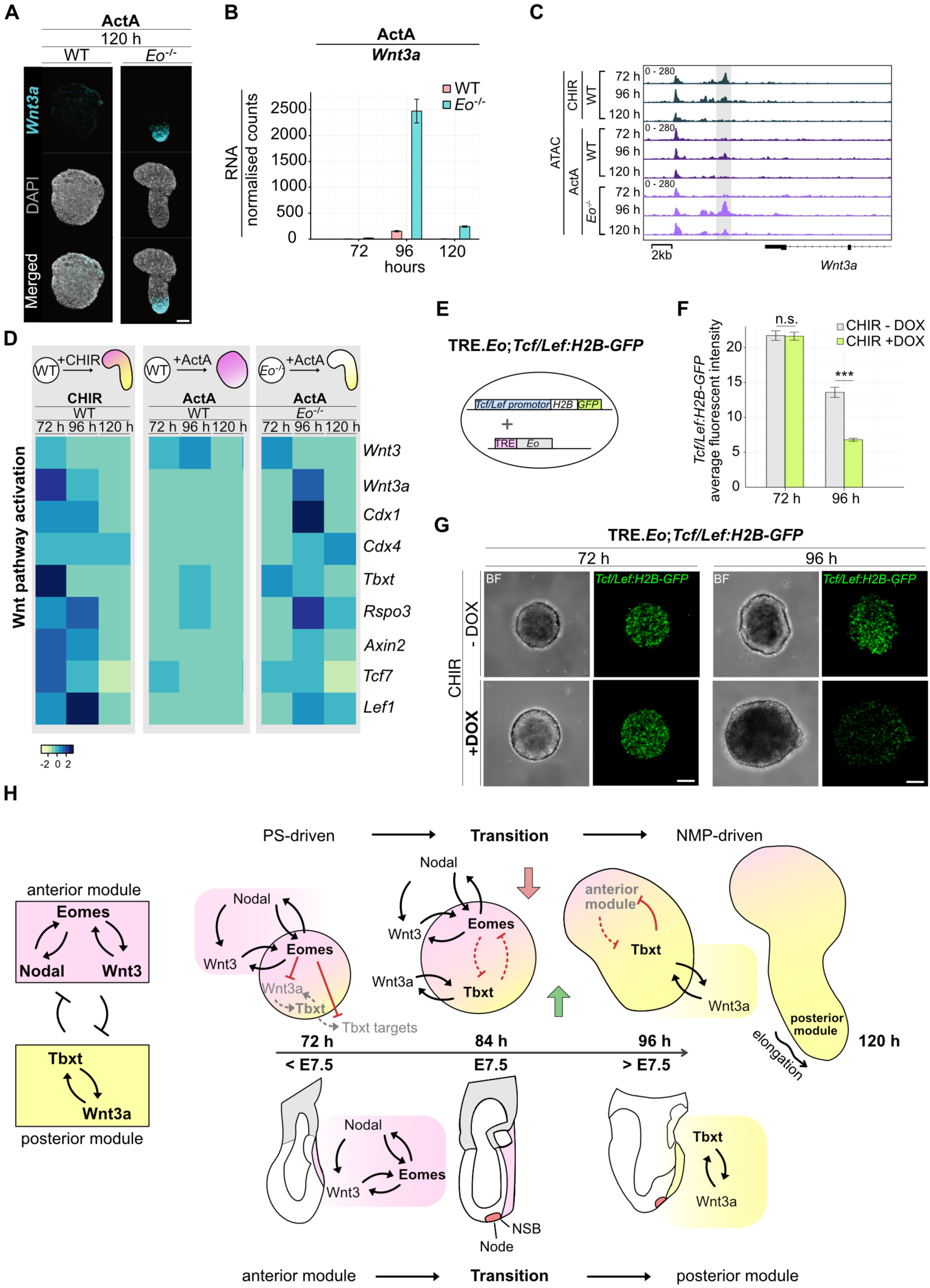
The canonical Wnt cascade of the posterior module is repressed at multiple levels by *Eomes*. **(A)** Fluorescent *in situ* hybridisation (FISH) and **(B)** mRNA expression levels of *Wnt3a* in ActA-induced WT and *Eo* ^−/−^ gastruloids, suggests suppression of *Wnt3a* by *Eomes*. ActA-induced *Eomes*-deficient (*Eo* ^−/−^) gastruloids show similar *Wnt3a* patterns compared to CHIR-induced WT (see Fig. 4A). n≥8. Scale bar 100 µm. **(C)** ATACseq coverage tracks at the gene locus of *Wnt3a* show accessible chromatin at a putative enhancer of CHIR-treated WT gastruloids, and reduced chromatin accessibility when WT gastruloids are induced with ActA. ActA-pulsed *Eo* ^−/−^ gastruloids, restore levels of chromatin accessibility. **(D)** Heatmap representation of expression of components of the canonical Wnt-signalling cascade. Heatmaps show normalized RNAseq expression data of indicated genes in CHIR- and ActA-treated WT gastruloids, and in ActA-treated *Eo* ^−/−^ gastruloids. CHIR-treated WT gastruloids exhibit an active Wnt signature (*Axin2*, *Tcf7*, *Lef1*), that is reduced in ActA-induced gastruloids. ActA-pulsed *Eo* ^−/−^ gastruloids establish an active Wnt- and tailbud signature (*Cdx1, Cdx4, Tbxt, Rspo3*). Gastruloid schematics above the heatmaps illustrate observed phenotypes and Wnt signature activities in different conditions. **(E)** Schematic of the *Eomes*-inducible (TRE.*Eo)* cell line that harbours the Tcf/Lef:H2B-GFP transcriptional Wnt-reporter. **(F)** Bar chart representation of Wnt-reporter activity in CHIR-pulsed gastruloids after forced expression of *Eomes.* Fluorescent intensities of the *H2B-GFP* reporter are compared between uninduced (-DOX) and *Eomes*-expressing (+DOX) gastruloids at 72 h and 96 h, demonstrating reduced Wnt-reporter activation in *Eomes-*expressing cells at 96 h (***, p<0,001). **(G)** Examples of brightfield and GFP-fluorescent images of uninduced and *Eomes*-expressing CHIR-pulsed gastruloids as used for reporter quantification in (F). n≥20. Scale bars 100 µm. **(H)** A comprehensive modal describing the interdependencies of *Nodal* and *Wnt* signals with Tbx TFs *Eomes* and *Tbxt,* that guide the progression of gastrulation from early primitive streak stages to axial elongation from progenitor pools. The anterior regulatory module (*Eomes*/*Nodal*/*Wnt3*) dominates the signalling landscape of the PS before E7.5 and represses activities of the posterior module (*Brachyury/ Wnt3a)* that promotes caudal axial elongation from NMPs. The timeline and schematics of embryonic stages indicate the corresponding timepoint of regulation in gastruloids.

To corroborate our finding that *Eomes* broadly impacts canonical Wnt signalling, we engineered a mESC cell line that harbours a Tcf/Lef:H2B-GFP canonical Wnt reporter cassette (Ferrer-Vaquer et al., 2010) and the DOX-inducible *Eomes* expression cassette (Fig. 6E; TRE:Eo; Tcf/Lef:H2B-GFP) and generated gastruloids using CHIR induction (Fig. 6F, G). We compared reporter expression in CHIR induced gastruloids in the presence and absence of forced *Eomes* expression. While at 72 h there was no major difference, at 96 h H2B-GFP reporter expression was significantly reduced after forced *Eomes* expression (Fig. 6F, G), confirming the observation of globally reduced canonical Wnt-signalling responses by transcriptional profiling (Fig. 6D).

In summary, we find that *Nodal*/*ActivinA*-signalling fully relies on *Eomes* to suppress activities of the posterior functional module of *Tbxt* together with canonical *Wnt3a* signalling for posterior axis elongation. Repression of the posterior module is achieved at multiple different levels, such as reduced expression of *Wnt3a* and *Tbxt* (Fig. 5A, 6A), repression of signal transduction of the canonical Wnt-cascade (Fig. 6F), and repression of *TBXT* chromatin functions (Fig. 4K, 5C) (Schüle et al., 2023).

## Discussion

Previous studies uncovered the core signals and transcriptional regulators that orchestrate the generation of cells that build the basic body plan in a correct spatial and temporal order. Here, we studied the underlying molecular regulation and interdependencies of the factors that pattern the gastrulation process of mammalian embryos. We combined genetic and pharmacological approaches in highly versatile mouse gastruloids to overcome the experimental limitations of mammalian development. Our experiments reveal the cross-regulation of two competing functional modules that control spatial patterning and the temporal progression during the first two days of gastrulation as represented in a comprehensive model (Fig. 6H). The consecutively acting regulatory modules guide the proper specification of different cell types that is accompanied by a transition of different morphogenetic modes for cell recruitment during early and late germ layer formation. Gastrulation is initiated when combined activities of core signals of the anterior module, *Nodal* and *Wnt3* induce *Eomes* expression that is crucial for the specification of early AM (including cranial, cardiovascular and anterior paraxial mesoderm) and DE from epiblast cells while they ingress through the PS. *Eomes* reciprocally maintains *Wnt3* and *Nodal* expression in a mutually reinforcing feed forward loop that dominates the signalling environment during the first day of gastrulation. The formation of the embryonic node at E7.5, marks a rapid change in the signalling environment by the abrupt loss of expression of the core components of the anterior functional module, *Nodal*, *Wnt3* and *Eomes*, when *Tbxt* and *Wnt3a* of the posterior functional module, gain functional competence. Both, *Tbxt* and *Wnt3a* are expressed before E7.5, but activities are suppressed by at least two mechanisms that largely rely on *Eomes* downstream of *Nodal*/Smad2/3 signalling: First, by direct inhibition of Tbxt target genes (Schüle et al., 2023) (Fig. 5I), and, second, by suppression of the canonical Wnt-signalling cascade through transcriptional regulation of pathway members (Fig. 6).

Surprisingly, irrespective of the high levels of homology of amino acid sequence between *Wnt* and *Wnt3a* (Roelink & Nusse, 1991), both ligands show substantially different functions during gastrulation. In the early PS *Wnt3* promotes epiblast cells to exit pluripotency, initiates *Tbxt* expression and contributes to the specification of early PS derivatives in the context of the anterior regulatory module (Dias et al., 2025) characterized by high *Nodal* signalling. In the late PS, *Wnt3* is replaced by *Wnt3a* to provide a dominant canonical Wnt signalling environment promoting NMP generation and axial elongation. To date it is unclear where these substantial functional differences originate from. These could result from the temporal context in which both ligands act, or from differences in levels of expression and/or signalling functions of both ligands.

Previous studies demonstrated key requirements of combined *Tbxt* and *Wnt3a* activities for the elongation of the posterior body axis (Martin & Kimelman, 2008), that is fuelled by cell recruitment from NMPs that initially form in vicinity of the node (Cambray & Wilson, 2007; Wymeersch et al., 2019). Presented lineage tracing (Fig. 1A) demonstrates that the mesoderm progenitors for the first 7-10 pairs of somites are generated before node formation (<E7.5) and thus under the influence of the anterior regulatory module from the PS. These findings align with phenotypes of *Tbxt* and *Wnt3a* mutant embryos that suggested different molecular requirements, independent of *TbxtWnt3a* for the generation of mesoderm of most anterior and of trunk and tail somites (Guibentif et al., 2021). The molecular switch from the anterior to the posterior regulatory module is rather abrupt and can be pinpointed to E7.5. It is currently not clear if the morphogenetic transition of cell recruitment from PS ingression to NMP-derived mesoderm also occurs in a rapid step, or if this reflects a gradual change in gastrulation modes (Mittnenzweig et al., 2021; Tzouanacou et al., 2009; Wilson et al., 2009; Wymeersch et al., 2021).

A major question that remains is how the abrupt breakdown of the anterior module is initiated, when the feed-forward loop of *Nodal*, *Wnt3* and *Eomes* suddenly breaks down. Classical CHIR-pulsed gastruloids in which the anterior program is initiated but rapidly overpowered by the posterior module suggests that Wnt3a signalling plays a role in this transition. In agreement, we find extended *Nodal*/*Wnt3/Eomes* expression in *Tbxt* ^−/−^ gastruloids (Fig. 4B). However, the rather rapid kinetics of the breakdown of the *Nodal*/*Wnt3/Eomes* feed-forward loop, and the presence of *Tbxt* and *Wnt3a* already before E7.5 strongly suggest additional mechanisms of regulation. These could include suppression of *Nodal*-signalling, e.g. by increased expression of NODAL-antagonists *Lefty1/2* or negative regulation of the critical NODAL co-receptor Cripto (*Tdgf1*), by transcriptional mechanisms such as rapid nuclear degradation of *Eomes*, or by rapid changes in the biophysical property of cells at the E7.5 PS that might impact on signalling as described previously (Brennan et al., 2001; Gsell et al., 2025; Pfeiffer et al., 2018). Based on the different levels of reciprocal regulation between the early and late acting module that we discovered in this study and that were partially also described previously (Ben-Haim et al., 2006; Martin & Kimelman, 2008; Pfeiffer et al., 2018; Yamaguchi et al., 1999), we assume that it is the sum of synergistically acting regulatory mechanisms that generate the highly robust patterns along the embryonic AP-body axis.

In addition to providing a framework for the integration of transcription factors with signals during mammalian gastrulation, this study provides an explanation for the variable and often low representation of anterior fates, in particular AM and DE, in classical gastruloids treated with a CHIR pulse. Suggested from the current study, and also confirmed by Dias and colleagues (Dias et al., 2025), an experimental increase of Nodal-signalling levels preceding the CHIR-pulse would allow to shift gastruloids towards the anterior cell types of AM and DE to resemble earlier phases of PS development and gastrulation in this model system.

Future studies will be required to integrate how additional cues, including biophysical properties of 3D model systems, contribute to the remarkable robustness of stereotypic embryonic patterning. This robustness is largely based on the formation of patterns as intrinsic feature of self-organizing biological systems. Interestingly, the formation of molecular patterns is independent of the underlying morphogenetic event since gastruloids lack the equivalent of a PS or the epiblast. Existing and future embryoid models will aim for adding back additional components that will allow to also recapitulate morphogenetic aspects of embryogenesis *in vitro*.

## Key messages

- The sequential switch between modes of gastrulation from the early primitive streak to stem cell-generated axial elongation is accompanied by changes in transcriptional and signalling signatures.
- *Eomes/Nodal/Wnt3* and *Brachyury/Wnt3a* act as two competing regulatory modules that coordinate the developmental progression from the primitive streak stage to axial elongation.
- The transition from early gastrulation to the formation of posterior trunk structures and axial elongation relies on the functional downregulation of the early *Eomes/Nodal/Wnt3* and activation of the later *Brachyury/Wnt3a* module.
- Functional modules show reciprocal regulation by feed-forward and feedback-regulation.

## Materials and Methods

### Cell lines and culture

E14-based A2lox mESCs were used throughout this study (Iacovino et al., 2014). *Tbxt*/*Bra ^−/−^* and *Eo ^−/−^* (single knockout) and double knockout (dKO) mESCs and A2lox mESCs harbouring dox-inducible expression cassettes for *Eomes*.GFP were described previously (Tosic et al., 2019). *Tcf*/*Lef*:*H2B*-*GFP* reporter cells were generated by transgenic integration of the linearized Tcf/Lef:H2B-GFP reporter plasmid (addgene, Plasmid #32610) (Ferrer-Vaquer et al., 2010), followed by screening for GFP-reporter expression in single clones. ESCs were transfected with 6 µg linearized vector by electroporation (Nucleofector ESC kit, Lonza), cultured on 0.1% gelatine-coated dishes in Dulbecco’s modified Eagle’s medium (DMEM) containing 15 % fetal bovine serum (FBS, Gibco), 2 mM L-glutamine, 1X non-essential amino acids (NEAA), 1 mM sodium-pyruvate, 1X penicillin/streptomycin (all from Gibco), 100 µM β-mercaptoethanol (Sigma), Leukemia inhibitory factor (ESGRO LIF, Merck Millipore, 1000 U/ml), and 2i: CHIR99021 (Axon Medchem, 1386, 3 µM) and PD0325901 (Axon Medchem, 1408, 1 µM). The medium was changed daily and mESCs passaged every other day.

### Generation of gastruloids

Gastruloids were generated according to standard protocols (van den Brink et al., 2020) as Matrigel-free version. In brief, mESCs were dissociated and resuspended in ESGRO Complete Basal Medium (Merck Millipore, SF002-500) and distributed as 300 cells per well into ultra-low attachment plates (Greiner, 650970). 48 h after seeding, the aggregates were pulsed with either 0.3 mM CHIR (Axon Medchem, 1386) or 25 ng/µl Activin A (R&D Systems, 338-AC) for 24 h. For induced gene expression from the A2lox locus, gastruloids were treated with 1µg/ml doxycycline (Sigma-Aldrich, D9891) simultaneous to the CHIR pulse. The medium was changed at 72 h and 96 h.

### Whole mount *in situ* hybridization

Whole-mount *in situ* hybridization (WISH) was performed according to standard protocols (Behringer, 2014). Probes and respective information can be requested from the authors. In brief, gastruloids were fixed in 4% PFA / PBS o/n at 4°C, dehydrated by a methanol series and stored in 100% methanol at −20°C. After rehydration gastruloids were bleached in 6 % H2O2 for 5 min, digested by 1.6 μg/ml ProteinaseK / PBT for 2 min, and postfixed in 4% PFA / 0.2 % glutaraldehyde for 20 min before prehybridization for 4 h and hybridization o/n according to standard protocols. DIG-labelled RNA probes are detected using anti-Digoxigenin-AP Fab fragments (Roche) in 1% sheep serum, 2% BBR in MAB (0.1 M Maleic acid, 0.3 M NaCl, NaOH, 1% Tween-20 in H2O, pH 7.5) and incubation at 4°C o/n. The antibody was washed by repeated washes in MAB (>24 h, RT), and colour reactions performed in BM purple staining solution (Roche) for 2-6 h at RT.

### Fluorescent *in situ* hybridization

For fluorescent *in situ* hybridization, we used HCR™ RNA-FISH probe sets, amplifiers and buffers according to the manufacturer’s protocols (Molecular instruments). In brief, gastruloids were collected, fixed in 4% PFA / PBS for 2 h, and stored in methanol as described for WISH. After rehydration, gastruloids were digested by 1.6 μg/ml Proteinase K / PBT for 2 min and postfixed in 4% PFA for 20 min before prehybridization for 30 min at 37°C. Hybridization was performed at 37°C in probe solution o/n. On the following day, gastruloids were repeatedly washed, incubated at RT in hairpin solution o/n, and washed with SSCT (5x sodium chloride sodium citrate, 0,1% Tween 20) before image acquisition.

### Immunofluorescence staining

Gastruloids were fixed in 4% PFA / PBS for 1 h at 4°C, permeabilized in 0.3% Triton X-100/ PBT for 30 min, and blocked in 1% BSA / PBT for 1h at RT. Primary antibody incubation was performed at 4°C o/n in 1% BSA / PBT, gastruloids washed 4x in PBT before secondary fluorescence-conjugated antibody incubation for 3 h followed by DAPI staining for 30 min at RT. Primary antibodies used were α-BRACHYURY (TBXT) (R&D Systems; AF2085) and α-EOMES (abcam; ab23345) at suggested dilutions. Secondary anti-goat and anti-rabbit Alexa Fluor antibodies conjugated with 488-, 546- or 647-flurophores (Thermo Fisher) were used at 1:1000 dilution.

### Imaging

Images were acquired on a Leica DMi8 Thunder Imager System or a Leica M165FC Stereo microscope. Images were processed in the Leica LASX software and by Affinity Photo. All brightfield and fluorescent images that were acquired with the Thunder microscope are 3D-maximum projections of z-stacks.

### RNAseq and analysis

For RNA sequencing, 12-48 gastruloids per replicate were used and total RNA isolated using the RNeasy Mini kit (Qiagen). Concentration of the isolated RNA was quantified using the NanoDrop (Thermo Fisher). Library preparation and sequencing was performed by Novogene Services, UK, from a minimum of three biological replicates for each condition.

FASTQ pre-processing was performed using the Galaxy biocomputing platform (Afgan et al., 2018). Low-quality bases and adapter-containing reads were trimmed using Trim Galore (Galaxy Version 0.4.3.1), and reads were aligned to the mouse reference genome mm10 (mm10_UCSC_07_15, RNA STAR71 Galaxy Version 2.7.2b) using default parameters. Duplicates were removed using RmDup (Galaxy version 2.0.1) and genomic features were counted using htseq-count (Galaxy Version 0.9.1) with the following settings: mode-union, stranded-no, minimum alignment quality-10 and feature type-exon. Differentially expressed genes (DEGs) were analysed using DESeq2 functions in R with filtering for adjusted p-value<0.05, log2(FC)>1.5 for upregulated genes and log2(FC)<-1.5 for downregulated genes comparing WT versus knockout gastruloids (*Tbxt* ^−/−^ or *Eo* ^−/−^) individually at each time point. Venn diagrams represent the overlap of up- and downregulated genes in *Tbxt ^−/−^* and *Eo ^−/−^* compared to WT in the respective condition. For GO term analysis, all DEGs for a genotype and signalling condition were analyzed using Enrichr (Xie et al., 2021). Terms from GO Biological Process 2023 are shown with their adjusted p-value that were computed using the Benjamini-Hochberg method and the total number of overlapping genes are indicated. Heatmaps were plotted with galaxy (version 23.1.2.dev0) using the heatmap2 tool. Prior, z-scores were calculated from the normalized counts using table compute. For heatmaps in Fig. 2 and Fig. 5, z-scores were computed row-wise from each Supplementary Table. CHIR and ActA conditions were computed separately and TRE.Eo gastruloids were computed in comparison to CHIR-treated WT gastruloids. For the heatmap representing components of Wnt pathway activation in Fig. 6D and Fig. S3C, z-scores were computed row-wise from all depicted condition. The gene list for heatmaps that depict the spatiotemporal progression of gastruloid development were generated semi-manually by selecting genes from clustered DEGs (CHIR, RNAseq data) that reflect the temporal progression in WT gastruloids. In addition, the list contains selected genes from GO terms pattern specification process (GO:0007389) and axis specification (GO:0009798). The same gene lists were used for all conditions (see Supplementary Tables 2-4). Boxplots showing RNA expression levels were generated using geom_bar of the ggplot2 package in R. RNA expression levels are plotted as normalised counts of three or four replicates. Error bars indicate the standard error of the mean (SEM).

### ATACseq and analysis

ATACseq was performed according to published protocols (Buenrostro et al., 2013) with modifications as outlined below. In brief, gastruloids at indicated time points were dissociated and washed. 100,000 cells were lysed in 50 µl ATAC-seq lysis buffer and tagmentation performed with 50 µl of transposition reaction mix containing Tagment DNA Enzyme 1 (Nextera DNA Library Preparation Kit, Illumina) at 37°C and 600 rpm for one hour. DNA was purified using the Qiagen MinElute Kit and amplified by PCR (12 - 13 cycles) using Custom Nextera PCR primer and the NEBNext Ultra II Q5 Master Mix (Nextera DNA Library Preparation Kit (New England BioLabs, M0544S). PCR products were purified (Qiagen MinElute Kit), DNA concentration of the libraries measured with a Qubit Fluorometer (Thermo Fischer, Q32854) and fragment size determined with a Bioanalyzer (Agilent). Libraries with large fragment sizes were size selected for fragments <1000 bp using SPRI select beads. Sequencing was performed by Novogene Services, UK. Low-quality bases and adapter-containing reads were trimmed using Galaxy platform Trim Galore (Galaxy Version 0.4.3.1). Reads were mapped to the mm10 genome using Galaxy platform Bowtie2 v2.3.4.3+galaxy0) (Langmead & Salzberg, 2012) with the paired end option and the duplicates removed with RmDup (Galaxy version 2.0.1). To generated IGV tracks, the coverage files were created using bamCoverage54 (v3.3.2.0.0) with bin size 10 bases and normalization to RPKM and paired-end extension of the fragment size.

### Data acquisition and statistics

For the statistical analysis of gastruloid phenotype, lengths and widths were measured using the LASX software (Leica) from a minimum of three replicate experiments and n≥20 gastruloids for each condition and replicate (Supplementary Table 1). The fluorescent intensity of *Tcf/Lef:H2B-GFP* reporter expression was quantified using ImageJ2 (version 2.14.0/154f). A ROI was defined in the brightfield image and the mean gray value in the GFP channel measured. We measured the fluorescent intensities for n≥20 gastruloids per condition and time point. The statistical significance between control and CHIR +DOX treatment was calculated using a two-tailed Welch’s t-test (unequal variances t-test). Box plots and bar graphs were plotted using ggplot2 in R.

### Visualization of scRNAseq data from the mouse gastrulation atlas data

We used published single-cell RNA-seq data of whole mouse embryos (Pijuan-Sala et al., 2019). We plotted UMAPs of genes of interest for the time points E7.0, E7.5 and E8.0. UMAPS and overviews of all cell types were downloaded from https://marionilab.cruk.cam.ac.uk/MouseGastrulation2018/ (accessed 09/24). UMAPs in svg format were edited in Affinity Designer as follows: cells originating from extraembryonic tissues were removed, (visceral endoderm, ExE endoderm, ExE ectoderm and parietal endoderm). All cells expressing the indicated genes of interest were plotted in one colour (irrespective of levels of log2 normalized counts).

## Data availability

Sequencing datasets of CHIR and ActA-treated gastruloids have been deposited in the Gene Expression Omnibus (GEO) under accession code GSE283355 (ATACseq) and GSE283356 (RNAseq). ChIPseq data were previously published and are accessible under GSE194192.

## Acknowledgments

We thank T. Bass for technical assistance and M. Kyba for the A2lox.Cre mESC line, the Freiburg Galaxy Team, Bioinformatics, University of Freiburg (Germany) funded by the Collaborative Research Centre 992 Medical Epigenetics (DFG grant SFB 992/1 2012) and the German Federal Ministry of Education and Research BMBF grant 031 A538A de.NBI-RBC. This study was supported by the German Research Foundation (DFG) through the Heisenberg Program (AR 732/3-1), project grant (AR 732/2-1), project A08 of CRC 992 (project ID 192904750) to SJA, and Germany’s Excellence Strategy (CIBSS – EXC-2189 – Project ID 390939984) to KMS and SJA. The project was supported by the Ministry of Science and Education of the State of Baden Württemberg to SJA. KMS is supported by the EQUIP Program for Medical Scientist, Faculty of Medicine, University Freiburg and by a CRC 992 MEDEP-Fellowship (project ID 192904750). KM is supported by the Medical Research Council as part of UK Research and Innovation (MCUP1201/23). AD is supported by an EMBO Postdoctoral Fellowship (ALTF 948-2022) and a Generalitat de Catalunya AGAUR Grant (2021 SGR 00175). AMA is funded by an ERC Advanced Grant (834580) and the Maria de Maeztu Program for Units of Excellence in R&D (CEX2018-000792-M).

## Author contributions

AEW, JKS, FE, CMS, LZ and KMS performed experiments. AEW generated gastruloids, performed ISH, IF, imaging and RNA isolation. JKS generated gastruloids, analysed phenotypes, performed IF, imaging and ATAC. MT, AEW and JKS performed bioinformatical analyses. FE generated ESC lines and gastruloids. CMS, LZ and KMS contributed to ESC line generation and interpretation of data. KM performed and KM and SJA analysed time-lapse imaging. AEW, SP and SJA, analysed and interpreted the data. AD and AMA shared unpublished data and discussed data interpretation. AEW, SP, AMA and SJA, produced figures, wrote and edited the manuscript with input from all authors. SJA conceived the study.

## SUPPLEMENTARY FIGURES

**Fig. S1. Extended phenotype analysis of anterior-posterior axis elongation in gastruloids**

**(A, B)** Schematics of induction protocols to generate gastruloids as models of primitive streak patterning and development by **(A)** pulsing with CHIR to mimic Wnt signalling or **(B)** induction with Activin A (ActA) to induce *Nodal*-signalling.

**(C)** Schematics of cell lines used for the generation of gastruloids. Single knockouts were generated by the insertion of fluorescent reporters into one allele (*Eomes*^GFP^ and *Tbxt*^Tomato^) and frameshift deletions on the second allele. dKO cells are deficient for both *Eomes* and *Tbxt*.

**(D)** Examples of brightfield images showing the phenotypic variation of WT, *Tbxt* ^−/−^, *Eo* ^−/−^, and dKO gastruloids at 120 h following induction with either CHIR or ActA.

**(E)** Morphometric measurement of gastruloids of different genotypes and induction protocols. Gastruloids were measured along the maximal length and width as illustrated. Boxplots show the mean length (red) and mean width (blue) for each genotype after a CHIR- or ActA-pulse. For every condition three independent replicates with each n ≥20 gastruloids were measured (values provided in Supplementary table 1).

**(F, G)** Bar graphs show the calculated length/width ratios of gastruloids of different phenotype and induction protocol as read-out for phenotypic elongation.

**Fig. S2. RNAseq analysis featuring the molecular differences of WT, *Tbxt* ^−/−^ and *Eo* ^−/−^ gastruloids**

**(A-B)** Venn diagrams showing gene numbers of up- and downregulated differentially expressed genes (DEGs) at 72, 96 and 120 h in *Tbxt*^/-^ and *Eo* ^−/−^ gastruloids compared to WT and in **(A)** CHIR, or **(B)** ActA condition. *Tbxt ^−/−^* gastruloids exhibit markedly increased numbers of DEGs after the CHIR-pulse, while *Eo* ^−/−^ gastruloids exhibit higher numbers of DEGs following an ActA-pulse.

**(C-D)** GO term analysis for biological processes of all DEGs in *Tbxt* ^−/−^ and *Eo* ^−/−^ gastruloids in **(C)** CHIR, or **(D)** ActA conditions indicates gene function in axis specification and patterning. The -log(10) adjusted p-value for each term are plotted and total numbers of overlapping genes are indicated.

**Fig. S3. Analysis of effectors and regulators of the Wnt pathway in CHIR and ActA treated WT, *Tbxt* ^−/−^ and *Eo* ^−/−^ gastruloids**

**(A)** ATACseq coverage tracks at the gene loci of *Tcf7l1, and Dkk1* in CHIR pulsed WT and ActA-pulsed WT, and *Eo* ^−/−^ gastruloids at 72, 96 and 120 h. Putative repressor (*Tcf7l1*) or enhancer (*Dkk1*) sites are indicated in grey. Dynamic accessibility at a putative repressor site at the *Tcf7l1* locus is induced by CHIR in WT gastruloids, and by ActA in *Eo* ^−/−^ gastruloids, but not in ActA-pulsed WT gastruloids. The negative regulator of Wnt-signalling *Dkk1* shows the opposite regulation at a putative enhancer element. Counts normalized to RPKM are indicated.

**(B)** Bar graphs showing RNA expression levels of *Tcf7l1* and *Dkk1* that are generally reduced in *Eo* ^−/−^ gastruloids.

**(C)** Heatmaps of RNA expression levels of genes indicative for active Wnt signalling (e.g. *Rspo3*, *Axin2*, *Tcf7*, *Lef1*) and posterior marker genes (*Cdx1, Cdx4, Tbxt, Wnt3a*) in CHIR, or ActA-pulsed WT, *Eo* ^−/−^, and *Tbxt* ^−/−^ gastruloids. *Tbxt* ^−/−^ gastruloids *s*how reduced Wnt- and posterior signatures. ActA-pulsed WT and *Tbxt* ^−/−^ gastruloids globally lack posterior and Wnt-signatures, that are partially restored in *Eo* ^−/−^(ActA) gastruloids.

## Supplementary Material

**Supplementary Table 1**

Measured length and width of gastruloids of different genotypes (WT, *Tbxt ^−/−^*, *Eo ^−/−^*, dKO) and signalling pulses (CHIR or ActA). The table shows the individual measurements for length, width and for length/width ratios. For each genotype and signalling pulse (CHIR or ActA) n≥20 gastruloids were measured in three independent experiments. The median of measured values is indicated for each replicate. Relates to Figure S1.

**Supplementary Table 2**

Temporal progression of anterior-to-posterior marker gene expression in CHIR-treated gastruloids. RNAseq data show averaged normalised counts of marker genes (n=97) for the three genotypes (WT, *Tbxt ^−/−^*, *Eo ^−/−^*) and three time points (72, 96 and 120 h). Relates to Figure 2.

**Supplementary Table 3**

Temporal progression of anterior-to-posterior marker gene expression in ActA-treated gastruloids. RNAseq data shows averaged normalised counts of marker genes (n=97) for the three genotypes (WT, *Tbxt ^−/−^*, *Eo ^−/−^*) and three time points (72, 96 and 120 h). The same marker genes are listed as for CHIR-treated gastruloids in Table S2. Relates to Figure 2.

**Supplementary Table 4**

Comparison of the spatiotemporal progression of anterior to posterior marker gene expression between gastruloids with induced EOMES-GFP expression (TRE.Eo, +DOX), uninduced controls (-DOX) and CHIR-treated WT gastruloids. The same marker genes are listed as in heatmaps in Fig. 2. Table shows average normalised counts of RNAseq data. Relates to Figure 5.

## References

Afgan, E., Baker, D., Batut, B., van den Beek, M., Bouvier, D., Cech, M., Chilton, J., Clements, D., Coraor, N., Grüning, B. A., Guerler, A., Hillman-Jackson, J., Hiltemann, S., Jalili, V., Rasche, H., Soranzo, N., Goecks, J., Taylor, J., Nekrutenko, A., & Blankenberg, D. (2018). The Galaxy platform for accessible, reproducible and collaborative biomedical analyses: 2018 update. Nucleic Acids Research, 46(W1), W537–W544. 10.1093/nar/gky379

Aires, R., Dias, A., & Mallo, M. (2018). Deconstructing the molecular mechanisms shaping the vertebrate body plan. Current Opinion in Cell Biology, 55, 81–86. 10.1016/j.ceb.2018.05.009

Amin, S., Neijts, R., Simmini, S., Van Rooijen, C., Tan, S. C., Kester, L., Van Oudenaarden, A., Creyghton, M. P., & Deschamps, J. (2016). Cdx and T Brachyury Co-activate Growth Signaling in the Embryonic Axial Progenitor Niche. Cell Reports, 17(12), 3165–3177. 10.1016/j.celrep.2016.11.069

Arnold, S. J., Hofmann, U. K., Bikoff, E. K., & Robertson, E. J. (2008). Pivotal roles for eomesodermin during axis formation,epithelium-to-mesenchyme transition and endoderm specification in the mouse. Development, 135(3), 501–511. 10.1242/dev.014357

Arnold, S. J., & Robertson, E. J. (2009). Making a commitment: Cell lineage allocation and axis patterning in the early mouse embryo. Nature Reviews Molecular Cell Biology, 10(2), 91–103. 10.1038/nrm2618

Arnold, S. J., Stappert, J., Bauer, A., Kispert, A., Herrmann, B. G., & Kemler, R. (2000). Brachyury is a target gene of the Wnt/beta-catenin signaling pathway. Mechanisms of Development, 91(1–2), 249–258. 10.1016/s0925-4773(99)00309-3

Aulehla, A., Wehrle, C., Brand-Saberi, B., Kemler, R., Gossler, A., Kanzler, B., & Herrmann, B. G. (2003). Wnt3a Plays a Major Role in the Segmentation Clock Controlling Somitogenesis. Developmental Cell, 4(3), 395–406. 10.1016/S1534-5807(03)00055-8

Bardot, E. S., & Hadjantonakis, A.-K. (2020). Mouse gastrulation: Coordination of tissue patterning, specification and diversification of cell fate. Mechanisms of Development, 163, 103617. 10.1016/j.mod.2020.103617

Beccari, L., Moris, N., Girgin, M., Turner, D. A., Baillie-Johnson, P., Cossy, A.-C., Lutolf, M. P., Duboule, D., & Arias, A. M. (2018). Multi-axial self-organization properties of mouse embryonic stem cells into gastruloids. Nature, 562(7726), 272–276. 10.1038/s41586-018-0578-0

Behringer, R. (with Gertsenstein, M., Nagy, K. V., & Nagy, A.). (2014). Manipulating the mouse embryo: A laboratory manual (Fourth edition.). Cold Spring Harbor Laboratory Press.

Bénazéraf, B., & Pourquié, O. (2013). Formation and Segmentation of the Vertebrate Body Axis. Annual Review of Cell and Developmental Biology, 29(1), 1–26. 10.1146/annurev-cellbio-101011-155703

Ben-Haim, N., Lu, C., Guzman-Ayala, M., Pescatore, L., Mesnard, D., Bischofberger, M., Naef, F., Robertson, E. J., & Constam, D. B. (2006). The Nodal Precursor Acting via Activin Receptors Induces Mesoderm by Maintaining a Source of Its Convertases and BMP4. Developmental Cell, 11(3), 313–323. 10.1016/j.devcel.2006.07.005

Brennan, J., Lu, C. C., Norris, D. P., Rodriguez, T. A., Beddington, R. S. P., & Robertson, E. J. (2001). Nodal signalling in the epiblast patterns the early mouse embryo. Nature, 411(6840), 965–969. 10.1038/35082103

Buenrostro, J. D., Giresi, P. G., Zaba, L. C., Chang, H. Y., & Greenleaf, W. J. (2013). Transposition of native chromatin for fast and sensitive epigenomic profiling of open chromatin, DNA-binding proteins and nucleosome position. 10.1038/nmeth.2688

Cambray, N., & Wilson, V. (2007). Two distinct sources for a population of maturing axial progenitors. Development, 134(15), 2829–2840. 10.1242/dev.02877

Collignon, J., Varlet, I., & Robertson, E. J. (1996). Relationship between asymmetric nodal expression and the direction of embryonic turning. Nature, 381(6578), 155–158. 10.1038/381155a0

Costello, I., Pimeisl, I.-M., Dräger, S., Bikoff, E. K., Robertson, E. J., & Arnold, S. J. (2011). The T-box transcription factor Eomesodermin acts upstream of Mesp1 to specify cardiac mesoderm during mouse gastrulation. Nature Cell Biology, 13(9), 1084–1091. 10.1038/ncb2304

Dias, A., Pascual-Mas, P., Torregrosa-Cortés, G., McNamara, H. M., Wehmeyer, A. E., Arnold, S. J., & Arias, A. M. (2025). A temporal coordination between Nodal and Wnt signalling governs the emergence of the mammalian body plan (p. 2025.01.11.632562). bioRxiv. 10.1101/2025.01.11.632562

Downs, K. M., & Davies, T. (1993). Staging of gastrulating mouse embryos by morphological landmarks in the dissecting microscope. Development, 118(4), 1255–1266. 10.1242/dev.118.4.1255

Duarte, P., Brattig Correia, R., Nóvoa, A., & Mallo, M. (2023). Regulatory changes associated with the head to trunk developmental transition. BMC Biology, 21(1),170. 10.1186/s12915-023-01675-2

Dunn, N. R., Vincent, S. D., Oxburgh, L., Robertson, E. J., & Bikoff, E. K. (2004). Combinatorial activities of Smad2 and Smad3 regulate mesoderm formation and patterning in the mouse embryo. Development, 131(8), 1717–1728. 10.1242/dev.01072

Evans, A. L., Faial, T., Gilchrist, M. J., Down, T., Vallier, L., Pedersen, R. A., Wardle, F. C., & Smith, J. C. (2012). Genomic Targets of Brachyury (T) in Differentiating Mouse Embryonic Stem Cells. PLoS ONE, 7(3). 10.1371/journal.pone.0033346

Ferrer-Vaquer, A., Piliszek, A., Tian, G., Aho, R. J., Dufort, D., & Hadjantonakis, A.-K. (2010). A sensitive and bright single-cell resolution live imaging reporter of Wnt/ß-catenin signaling in the mouse. 10.1186/1471-213X-10-121

Gsell, S., Tlili, S., Merkel, M., & Lenne, P.-F. (2025). Marangoni-like tissue flows enhance symmetry breaking of embryonic organoids. Nature Physics, 21(4), 644–653. 10.1038/s41567-025-02802-2

Guibentif, C., Griffiths, J. A., Imaz-Rosshandler, I., Ghazanfar, S., Nichols, J., Wilson, V., Göttgens, B., & Marioni, J. C. (2021). Diverse Routes toward Early Somites in the Mouse Embryo. Developmental Cell, 56(1), 141–153.e6. 10.1016/j.devcel.2020.11.013

Herrmann, B. G., Labeit, S., Poustka, A., King, T. R., & Lehrach, H. (1990). Cloning of the T gene required in mesoderm formation in the mouse. Nature, 343(6259), 617–622. 10.1038/343617a0

Iacovino, M., Roth, M. E., & Kyba, M. (2014). Rapid Genetic Modification of Mouse Embryonic Stem Cells by Inducible Cassette Exchange Recombination. In M. F. Ochs (Ed.), Gene Function Analysis (Vol. 1101, pp. 339–351). Humana Press. 10.1007/978-1-62703-721-1_16

Kinder, S. J., Tsang, T. E., Quinlan, G. A., Hadjantonakis, A.-K., Nagy, A., & Tam, P. P. L. (1999). The orderly allocation of mesodermal cells to the extraembryonic structures and the anteroposterior axis during gastrulation of the mouse embryo. Development, 126(21), 4691–4701. 10.1242/dev.126.21.4691

Kispert, A., & Herrmann, B. G. (1994). Immunohistochemical Analysis of the *Brachyury* Protein in Wild-Type and Mutant Mouse Embryos. Developmental Biology, 161(1), 179–193. 10.1006/dbio.1994.1019

Koch, F., Scholze, M., Wittler, L., Schifferl, D., Sudheer, S., Grote, P., Timmermann, B., Macura, K., & Herrmann, B. G. (2017). Antagonistic Activities of Sox2 and Brachyury Control the Fate Choice of Neuro-Mesodermal Progenitors. Developmental Cell, 42(5), 514–526.e7. 10.1016/j.devcel.2017.07.021

Kwon, G. S., Viotti, M., & Hadjantonakis, A.-K. (2008). The Endoderm of the Mouse Embryo Arises by Dynamic Widespread Intercalation of Embryonic and Extraembryonic Lineages. Developmental Cell, 15(4), 509–520. 10.1016/j.devcel.2008.07.017

Langmead, B., & Salzberg, S. L. (2012). Fast gapped-read alignment with Bowtie 2. Nature Methods, 9(4), 357–359. 10.1038/nmeth.1923

Lawson, K. A., Meneses, J. J., & Pedersen, R. A. (1986). Cell fate and cell lineage in the endoderm of the presomite mouse embryo, studied with an intracellular tracer. Developmental Biology, 115(2), 325–339. 10.1016/0012-1606(86)90253-8

Liu, P., Wakamiya, M., Shea, M. J., Albrecht, U., Behringer, R. R., & Bradley, A. (1999). Requirement for Wnt3 in vertebrate axis formation. Nature Genetics, 22(4), 361–365. 10.1038/11932

Lu, C. C., & Robertson, E. J. (2004). Multiple roles for Nodal in the epiblast of the mouse embryo in the establishment of anterior-posterior patterning. Developmental Biology, 273(1), 149–159. 10.1016/j.ydbio.2004.06.004

Martin, B. L., & Kimelman, D. (2008). Regulation of Canonical Wnt Signaling by Brachyury Is Essential for Posterior Mesoderm Formation. Developmental Cell, 15(1), 121–133. 10.1016/j.devcel.2008.04.013

McDole, K., Guignard, L., Amat, F., Berger, A., Malandain, G., Royer, L. A., Turaga, S. C., Branson, K., & Keller, P. J. (2018). In Toto Imaging and Reconstruction of Post-Implantation Mouse Development at the Single-Cell Level. Cell, 175(3), 859–876.e33. 10.1016/j.cell.2018.09.031

McNamara, H. M., Solley, S. C., Adamson, B., Chan, M. M., & Toettcher, J. E. (2024). Recording morphogen signals reveals mechanisms underlying gastruloid symmetry breaking. Nature Cell Biology, 26(11), 1832–1844. 10.1038/s41556-024-01521-9

Mittnenzweig, M., Mayshar, Y., Cheng, S., Ben-Yair, R., Hadas, R., Rais, Y., Chomsky, E., Reines, N., Uzonyi, A., Lumerman, L., Lifshitz, A., Mukamel, Z., Orenbuch, A.-H., Tanay, A., & Stelzer, Y. (2021). A single-embryo, single-cell time-resolved model for mouse gastrulation. Cell, 184(11), 2825–2842.e22. 10.1016/j.cell.2021.04.004

Morgani, S. M., & Hadjantonakis, A.-K. (2020). Signaling regulation during gastrulation: Insights from mouse embryos and in vitro systems. In Current Topics in Developmental Biology (Vol. 137, pp. 391–431). Elsevier. 10.1016/bs.ctdb.2019.11.011

Norris, D. P., Brennan, J., Bikoff, E. K., & Robertson, E. J. (2002). The Foxh1-dependent autoregulatory enhancer controls the level of Nodal signals in the mouse embryo. Development, 129(14), 3455–3468. 10.1242/dev.129.14.3455

Norris, D. P., & Robertson, E. J. (1999). Asymmetric and node-specific nodal expression patterns are controlled by two distinct cis-acting regulatory elements. Genes & Development, 13(12), 1575–1588. 10.1101/gad.13.12.1575

Pfeiffer, M. J., Quaranta, R., Piccini, I., Fell, J., Rao, J., Röpke, A., Seebohm, G., & Greber, B. (2018). Cardiogenic programming of human pluripotent stem cells by dose-controlled activation of EOMES. Nature Communications, 9(1), 440. 10.1038/s41467-017-02812-6

Pijuan-Sala, B., Griffiths, J. A., Guibentif, C., Hiscock, T. W., Jawaid, W., Calero-Nieto, F. J., Mulas, C., Ibarra-Soria, X., Tyser, R. C. V., Ho, D. L. L., Reik, W., Srinivas, S., Simons, B. D., Nichols, J., Marioni, J. C., & Göttgens, B. (2019). A single-cell molecular map of mouse gastrulation and early organogenesis. Nature, 566(7745), 490–495. 10.1038/s41586-019-0933-9

Probst, S., Sagar, Tosic, J., Schwan, C., Grün, D., & Arnold, S. J. (2021). Spatiotemporal sequence of mesoderm and endoderm lineage segregation during mouse gastrulation. Development, 148(1), dev193789. 10.1242/dev.193789

Rivera-Pérez, J. A., & Magnuson, T. (2005). Primitive streak formation in mice is preceded by localized activation of Brachyury and Wnt3. Developmental Biology, 288(2), 363–371. 10.1016/j.ydbio.2005.09.012

Robertson, E. J. (2014). Dose-dependent Nodal/Smad signals pattern the early mouse embryo. Seminars in Cell & Developmental Biology, 32, 73–79. 10.1016/j.semcdb.2014.03.028

Roelink, H., & Nusse, R. (1991). Expression of two members of the Wnt family during mouse development—Restricted temporal and spatial patterns in the developing neural tube. Genes & Development, 5(3), 381–388. 10.1101/gad.5.3.381

Rossi, G., Broguiere, N., Miyamoto, M., Boni, A., Guiet, R., Girgin, M., Kelly, R. G., Kwon, C., & Lutolf, M. P. (2021). Capturing Cardiogenesis in Gastruloids. Cell Stem Cell, 28(2), 230–240.e6. 10.1016/j.stem.2020.10.013

Savory, J. G. A., Bouchard, N., Pierre, V., Rijli, F. M., De Repentigny, Y., Kothary, R., & Lohnes, D. (2009). Cdx2 regulation of posterior development through non-Hox targets. Development, 136(24), 4099–4110. 10.1242/dev.041582

Schröder, C. M., Zissel, L., Mersiowsky, S.-L., Tekman, M., Probst, S., Schüle, K. M., Preissl, S., Schilling, O., Timmers, H. Th. M., & Arnold, S. J. (2025). EOMES establishes mesoderm and endoderm differentiation potential through SWI/SNF-mediated global enhancer remodeling. Developmental Cell, 60(5), 735–748.e5. 10.1016/j.devcel.2024.11.014

Schüle, K. M., Weckerle, J., Probst, S., Wehmeyer, A. E., Zissel, L., Schröder, C. M., Tekman, M., Kim, G.-J., Schlägl, I.-M., Sagar, & Arnold, S. J. (2023). Eomes restricts Brachyury functions at the onset of mouse gastrulation. Developmental Cell, 58(18), 1627–1642.e7. 10.1016/j.devcel.2023.07.023

Simon, C. S., Downes, D. J., Gosden, M. E., Telenius, J., Higgs, D. R., Hughes, J. R., Costello, I., Bikoff, E. K., & Robertson, E. J. (2017). Functional characterisation of cis-regulatory elements governing dynamic Eomes expression in the early mouse embryo. Development, 144(7), 1249–1260. 10.1242/dev.147322

Suppinger, S., Zinner, M., Aizarani, N., Lukonin, I., Ortiz, R., Azzi, C., Stadler, M. B., Vianello, S., Palla, G., Kohler, H., Mayran, A., Lutolf, M. P., & Liberali, P. (2023). Multimodal characterization of murine gastruloid development. Cell Stem Cell, 30(6), 867–884.e11. 10.1016/j.stem.2023.04.018

Takada, S., Stark, K. L., Shea, M. J., Vassileva, G., McMahon, J. A., & McMahon, A. P. (1994). Wnt-3a regulates somite and tailbud formation in the mouse embryo. Genes & Development, 8(2), 174–189. 10.1101/gad.8.2.174

Ten Berge, D., Koole, W., Fuerer, C., Fish, M., Eroglu, E., & Nusse, R. (2008). Wnt Signaling Mediates Self-Organization and Axis Formation in Embryoid Bodies. Cell Stem Cell, 3(5), 508–518. 10.1016/j.stem.2008.09.013

Teo, A. K. K., Arnold, S. J., Trotter, M. W. B., Brown, S., Ang, L. T., Chng, Z., Robertson, E. J., Dunn, N. R., & Vallier, L. (2011). Pluripotency factors regulate definitive endoderm specification through eomesodermin. Genes & Development, 25(3), 238–250. 10.1101/gad.607311

Tosic, J., Kim, G.-J., Pavlovic, M., Schröder, C. M., Mersiowsky, S.-L., Barg, M., Hofherr, A., Probst, S., Köttgen, M., Hein, L., & Arnold, S. J. (2019). Eomes and Brachyury control pluripotency exit and germ-layer segregation by changing the chromatin state. Nature Cell Biology, 21(12), 1518–1531. 10.1038/s41556-019-0423-1

Turner, D. A., Girgin, M., Alonso-Crisostomo, L., Trivedi, V., Baillie-Johnson, P., Glodowski, C. R., Hayward, P. C., Collignon, J., Gustavsen, C., Serup, P., Steventon, B., Lutolf, M., & Martinez, A. A. (2017). Anteroposterior polarity and elongation in the absence of extraembryonic tissues and spatially localised signalling in *Gastruloids*, mammalian embryonic organoids. Development, dev.150391. 10.1242/dev.150391

Tzouanacou, E., Wegener, A., Wymeersch, F. J., Wilson, V., & Nicolas, J.-F. (2009). Redefining the Progression of Lineage Segregations during Mammalian Embryogenesis by Clonal Analysis. Developmental Cell, 17(3), 365–376. 10.1016/j.devcel.2009.08.002

Vallier, L., Mendjan, S., Brown, S., Chng, Z., Teo, A., Smithers, L. E., Trotter, M. W. B., Cho, C. H.-H., Martinez, A., Rugg-Gunn, P., Brons, G., & Pedersen, R. A. (2009). Activin/Nodal signalling maintains pluripotency by controlling Nanog expression. Development, 136(8), 1339–1349. 10.1242/dev.033951

van den Brink, S. C., Baillie-Johnson, P., Balayo, T., Hadjantonakis, A.-K., Nowotschin, S., Turner, D. A., & Martinez Arias, A. (2014). Symmetry breaking, germ layer specification and axial organisation in aggregates of mouse embryonic stem cells. Development, 141(22), 4231–4242. 10.1242/dev.113001

Veenvliet, J. V., Bolondi, A., Kretzmer, H., Haut, L., Scholze-Wittler, M., Schifferl, D., Koch, F., Guignard, L., Kumar, A. S., Pustet, M., Heimann, S., Buschow, R., Wittler, L., Timmermann, B., Meissner, A., & Herrmann, B. G. (2020). Mouse embryonic stem cells self-organize into trunk-like structures with neural tube and somites. Science, 370(6522), eaba4937. 10.1126/science.aba4937

Vincent, S. D., Dunn, N. R., Hayashi, S., Norris, D. P., & Robertson, E. J. (2003). Cell fate decisions within the mouse organizer are governed by graded Nodal signals. Genes & Development, 17(13), 1646–1662. 10.1101/gad.1100503

Viotti, M., Foley, A. C., & Hadjantonakis, A.-K. (2014). Gutsy moves in mice: Cellular and molecular dynamics of endoderm morphogenesis. Philosophical Transactions of the Royal Society B: Biological Sciences, 369(1657), 20130547. 10.1098/rstb.2013.0547

Wang, J., Sinha, T., & Wynshaw-Boris, A. (2012). Wnt Signaling in Mammalian Development: Lessons from Mouse Genetics. Cold Spring Harbor Perspectives in Biology, 4(5), a007963–a007963. 10.1101/cshperspect.a007963

Wehmeyer, A. E., Schüle, K. M., Conrad, A., Schröder, C. M., Probst, S., & Arnold, S. J. (2022). Chimeric 3D gastruloids – a versatile tool for studies of mammalian peri-gastrulation development. Development, 149(22), dev200812. 10.1242/dev.200812

Williams, M., Burdsal, C., Periasamy, A., Lewandoski, M., & Sutherland, A. (2012). Mouse primitive streak forms in situ by initiation of epithelial to mesenchymal transition without migration of a cell population. Developmental Dynamics, 241(2), 270–283. 10.1002/dvdy.23711

Wilson, V., Olivera-Martinez, I., & Storey, K. G. (2009). Stem cells, signals and vertebrate body axis extension. Development, 136(10), 1591–1604. 10.1242/dev.021246

Wymeersch, F. J., Skylaki, S., Huang, Y., Watson, J. A., Economou, C., Marek-Johnston, C., Tomlinson, S. R., & Wilson, V. (2019). Transcriptionally dynamic progenitor populations organised around a stable niche drive axial patterning. *Development (Cambridge*, England), 146(1), dev168161. 10.1242/dev.168161

Wymeersch, F. J., Wilson, V., & Tsakiridis, A. (2021). Understanding axial progenitor biology *in vivo* and *in vitro*. Development, 148(4), dev180612. 10.1242/dev.180612

Xie, Z., Bailey, A., Kuleshov, M. V., Clarke, D. J. B., Evangelista, J. E., Jenkins, S. L., Lachmann, A., Wojciechowicz, M. L., Kropiwnicki, E., Jagodnik, K. M., Jeon, M., & Ma’ayan, A. (2021). Gene Set Knowledge Discovery with Enrichr. Current Protocols, 1(3), e90. 10.1002/cpz1.90

Yamaguchi, T. P. (2008). Genetics of Wnt Signaling During Early Mammalian Development. In E. Vincan (Ed.), Wnt Signaling (Vol. 468, pp. 287–305). Humana Press. 10.1007/978-1-59745-249-6_23

Yamaguchi, T. P., Takada, S., Yoshikawa, Y., Wu, N., & McMahon, A. P. (1999). T (Brachyury) is a direct target of Wnt3a during paraxial mesoderm specification. Genes & Development, 13(24), 3185–3190. 10.1101/gad.13.24.3185

Young, T., Rowland, J. E., Van De Ven, C., Bialecka, M., Novoa, A., Carapuco, M., Van Nes, J., De Graaff, W., Duluc, I., Freund, J.-N., Beck, F., Mallo, M., & Deschamps, J. (2009). Cdx and Hox Genes Differentially Regulate Posterior Axial Growth in Mammalian Embryos. Developmental Cell, 17(4), 516–526. 10.1016/j.devcel.2009.08.010

