## Supplementary figures and images for "Competing regulatory modules control the transition between mammalian gastrulation modes"

### Supplementary Figures

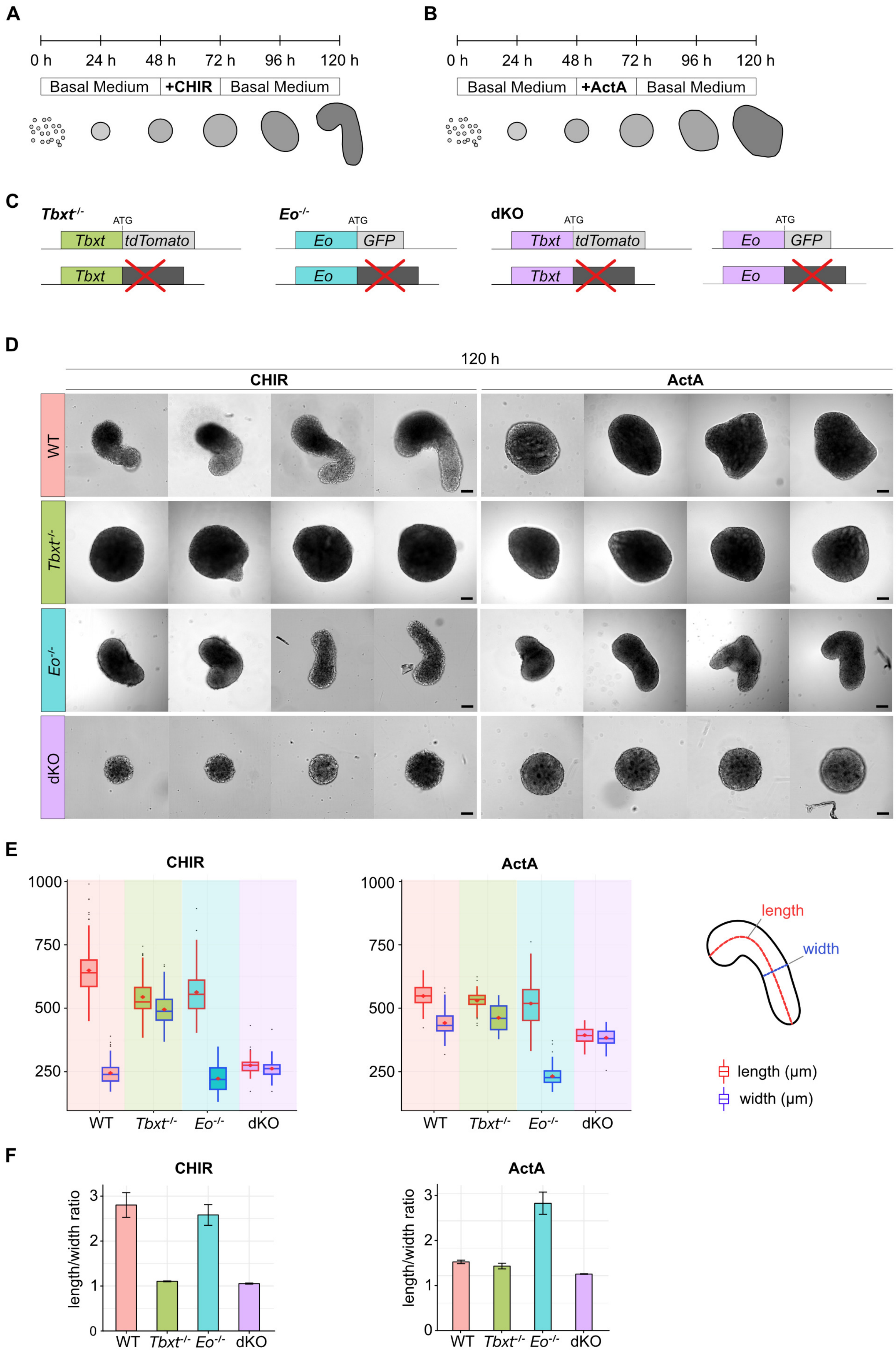

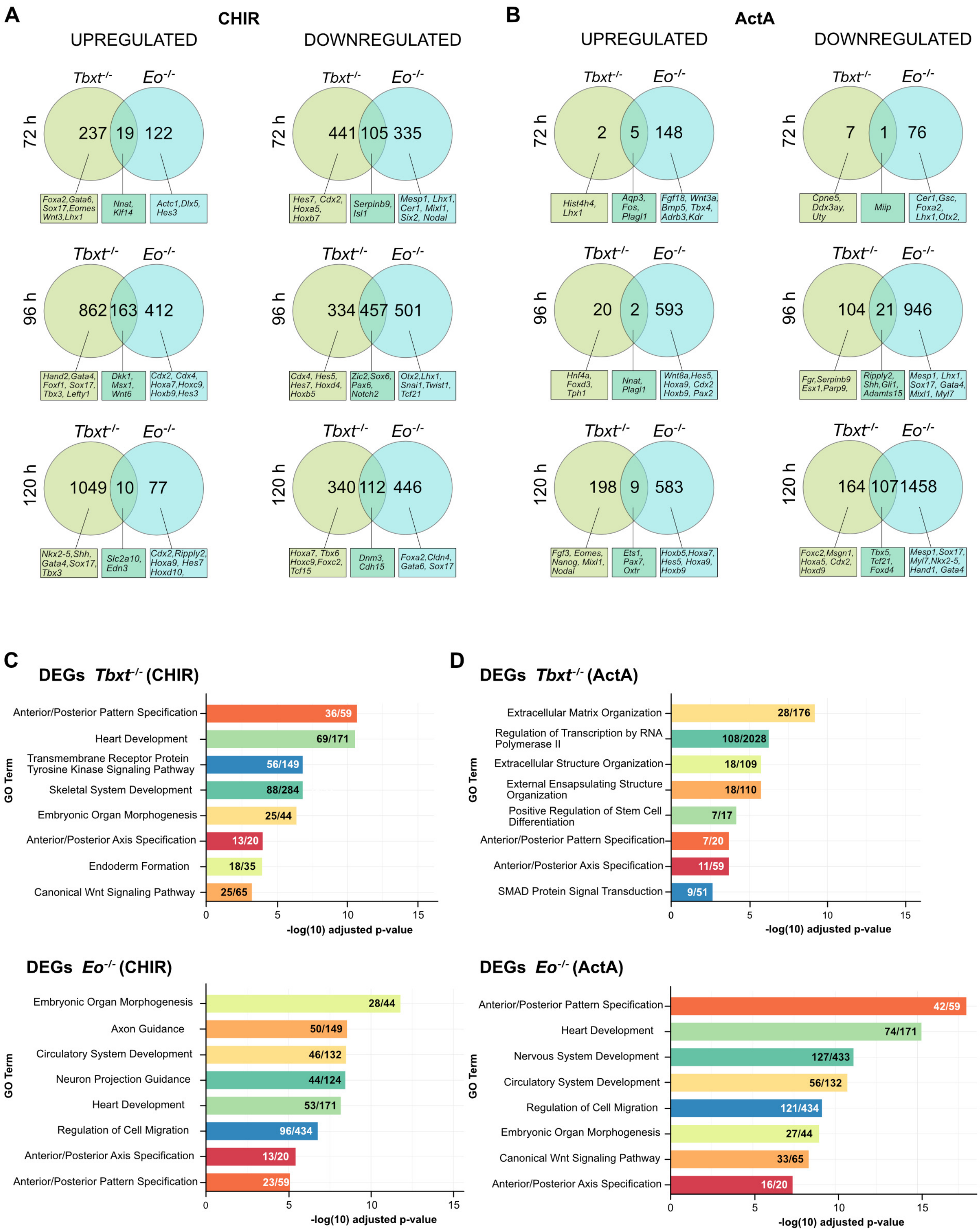

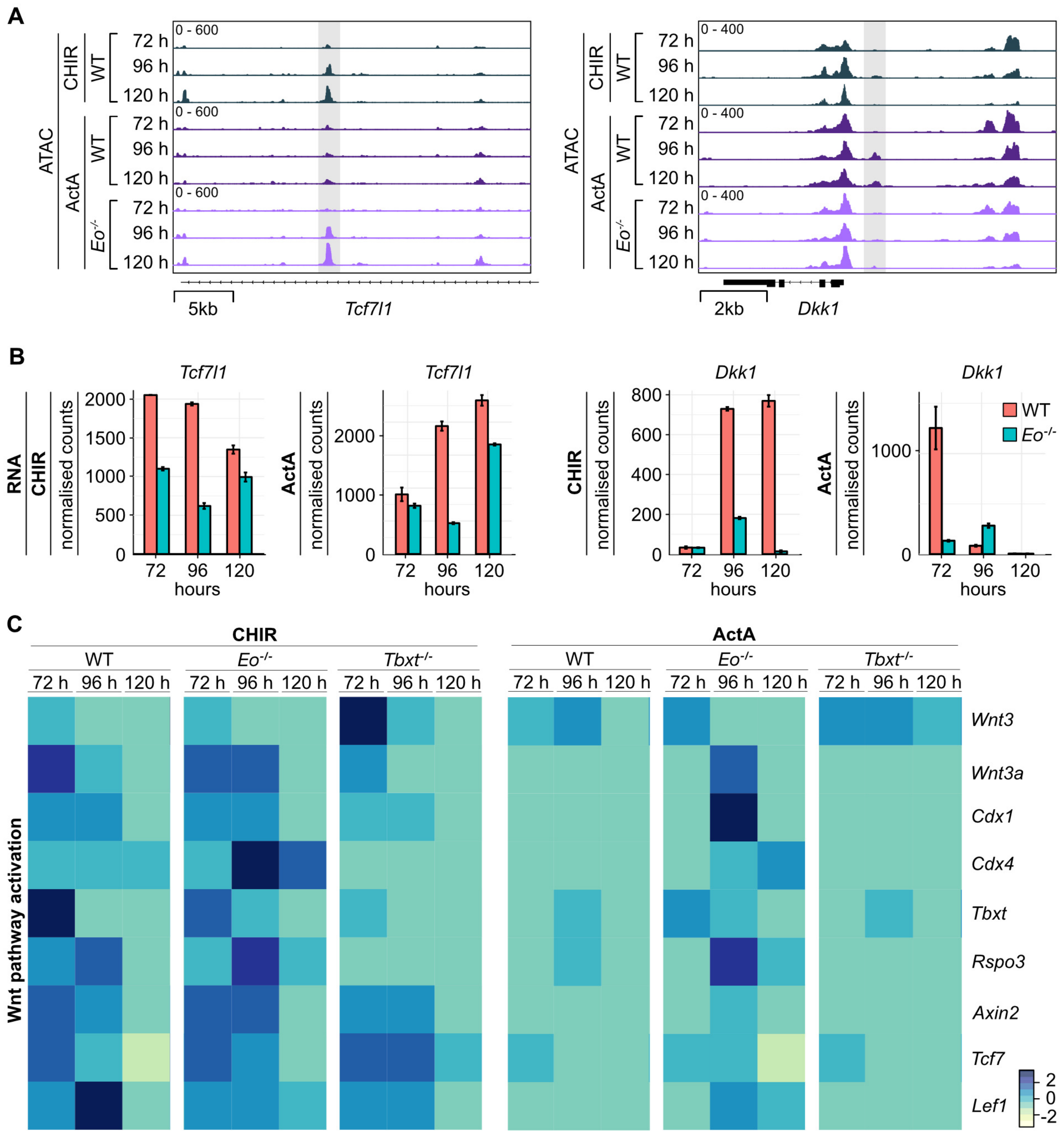
